# Cryo-EM structure of the human L-type amino acid transporter 1 in complex with glycoprotein CD98hc

**DOI:** 10.1101/577551

**Authors:** Yongchan Lee, Pattama Wiriyasermkul, Chunhuan Jin, Lili Quan, Ryuichi Ohgaki, Suguru Okuda, Tsukasa Kusakizako, Tomohiro Nishizawa, Kazumasa Oda, Ryuichiro Ishitani, Takeshi Yokoyama, Takanori Nakane, Mikako Shirouzu, Hitoshi Endou, Shushi Nagamori, Yoshikatsu Kanai, Osamu Nureki

## Abstract

The L-type amino acid transporter 1 (LAT1) transports large neutral amino acids and drugs across the plasma membrane and is crucial for nutrient uptake, brain drug delivery and tumor growth. LAT1 is a unique solute carrier that forms a disulfide-linked heterodimer with the cell-surface glycoprotein CD98 heavy chain (CD98hc), but the mechanisms of its molecular assembly and amino acid transport are poorly understood. Here we report the cryo-EM structure of the human LAT1-CD98hc heterodimer at 3.4 Å resolution, revealing the hitherto unprecedented architecture of a solute carrier-glycoprotein heterocomplex. LAT1 features a canonical LeuT-fold while exhibiting an unusual loop structure on transmembrane helix 6, creating an extended cavity to accommodate bulky hydrophobic amino acids and drugs. CD98hc engages with LAT1 through multiple interactions, not only in the extracellular and transmembrane domains but also in the interdomain linker. The heterodimer interface features multiple sterol molecules, corroborating previous biochemical data on the role of cholesterols in heterodimer stabilization. We also visualized the binding modes of two anti-CD98 antibodies and show that they recognize distinct, multiple epitopes on CD98hc but not its glycans, explaining their robust reactivities despite the glycan heterogeneity. Furthermore, we mapped disease-causing mutations onto the structure and homology models, which rationalized some of the phenotypes of SLC3- and SLC7-related congenital disorders. Together, these results shed light on the principles of the structural assembly between a glycoprotein and a solute carrier, and provide a template for improving preclinical drugs and therapeutic antibodies targeting LAT1 and CD98.

## Introduction

Amino acid transporters are key players in cellular metabolism. Humans possess more than 65 amino acid transporters, which belong to 11 discrete solute carrier (SLC) families *(1)*. The heteromeric amino acid transporters (HATs or SLC3/SLC7 families) are unique among all SLC transporters, in that they are composed of two subunits, the light chain (SLC7) and the heavy chain (SLC3), linked by a conserved disulfide-bridge *(2–4).* The light chain is a twelve-spanning membrane protein belonging to the amino acid-polyamine-organocation transporter (APC) superfamily *(5)* and functions as the catalytic subunit of the HATs *(6)*, whereas the heavy chain is a single-spanning type-II membrane glycoprotein that facilitates the heterodimer trafficking *(7)*. The human genome contains eight SLC7 members (SLC7A5–11, 13), which specifically associate with one of two SLC3 members (SLC3A1 and 2), giving rise to eight HAT subtypes *(8)*. These HATs show different tissue expression, localization, Na^+^-dependence and substrate specificity, and thereby play diverse roles in human physiology *(9)*.

The L-type amino acid transporter 1 (LAT1 or SLC7A5) is a Na^+^-independent amino acid antiporter with broad substrate specificity towards large neutral amino acids such as Leu, Tyr and Trp *(10).* LAT1 is highly expressed in immune T cells *(2, 11)*, the blood-brain barrier *(12)* and various types of cancers *(10, 13)*, and thus is an important target for cancer diagnosis *(14)* and treatment *(15, 16)*, Parkinson’s disease *(17)*, and the development of novel blood-brain barrier-crossing drugs *(18)*. In addition, Leu uptake by LAT1 is essential for mTOR activation *(19).* LAT1 also transports amino-acid derivatives, such as thyroid hormones *(20)*, L-DOPA (21), melphalan *(22)* and gabapentine *(22).* LAT1 forms a disulfide-linked heterodimer with CD98 heavy chain (CD98hc, also known as 4F2hc or SLC3A2) *(2).* The disulfide bridge is formed between two Cys residues located in the second extracellular loop of LAT1 and the linker connecting the extracellular and transmembrane domains of CD98hc *(23).* Apart from amino acid transport, CD98hc is involved in many cellular functions, ranging from cell adhesion *(24)* to immune activation *(25)* and the cellular entry of pathogens *(26, 27)*. In particular, CD98hc is a well-known interaction partner for integrins in mediating intracellular signaling *(28, 29)*.

Despite its biological importance, our understanding of the molecular assembly and function of LAT1-CD98hc is limited. Previous studies on the bacterial SLC7 homologs, ApcT *(30)* and AdiC *(31, 32)*, which lack the heavy chain, have provided the basis to understand the amino acid transport mechanism by the light chain. In addition, the crystal structure of an isolated extracellular domain of CD98hc has revealed its glucosidase-like fold *(33).* However, many questions remain unanswered. How do the heavy chain and the light chain interact with each other? How can LAT1 recognize a broad range of substrates, while its bacterial homologs cannot? What is the functional role of the single transmembrane domain of CD98hc? How does the heavy chain help the trafficking of the light chain, and how does the light chain prevent the degradation of the heavy chain *(34)*? What are the structure-function relationships of SLC3- and SLC7-related genetic diseases *(35–39)?* These questions can only be answered by studying the mammalian LAT1-CD98hc complex in its full heterodimeric assembly.

In this study, we used cryo-electron microscopy to reveal the molecular architecture and mechanisms of the human LAT1-CD98hc complex. Based on the high-resolution structures of LAT1-CD98hc bound to one or two monoclonal antibodies, we defined the mechanisms for heterodimer assembly, ligand recognition, disease-causing mutations and antibody targeting of LAT1-CD98hc. These results provide a framework to address many of the unanswered questions about this unique solute carrier-glycoprotein complex.

## Results

### Purification of the heterodimer

A major obstacle in the structural and biochemical analyses of the HATs was obtaining a natively-folded, disulfide-linked heterodimer. Although previous studies used *Escherichia coli (40)* or *Pichia pastoris (41)* for overexpression, we reasoned that the mammalian-type protein biogenesis pathway would be essential for reliable heterodimer production. Therefore, we screened a panel of constructs in mammalian cells and found that the co-expression of the N-terminally GFP-tagged full-length CD98hc isoform c and the C-terminally Flag epitope-tagged full-length LAT1 results in robust heterodimer formation (Fig. S1). We overproduced this full-length LAT1-CD98hc construct in HEK293S GnTI^−^ cells and purified the heterodimer through two rounds of antibody-based affinity purification (Fig. 1a and Fig. S1, also see Methods). The use of two cholesterol analogs, cholesteryl hemisuccinate and digitonin, markedly improved the stability of the heterodimer, consistent with a previous study *(42).* An SDS-PAGE analysis of the final product showed almost 100% formation of the disulfide-linked heterodimer (Fig. 1b).

**Figure 1.**
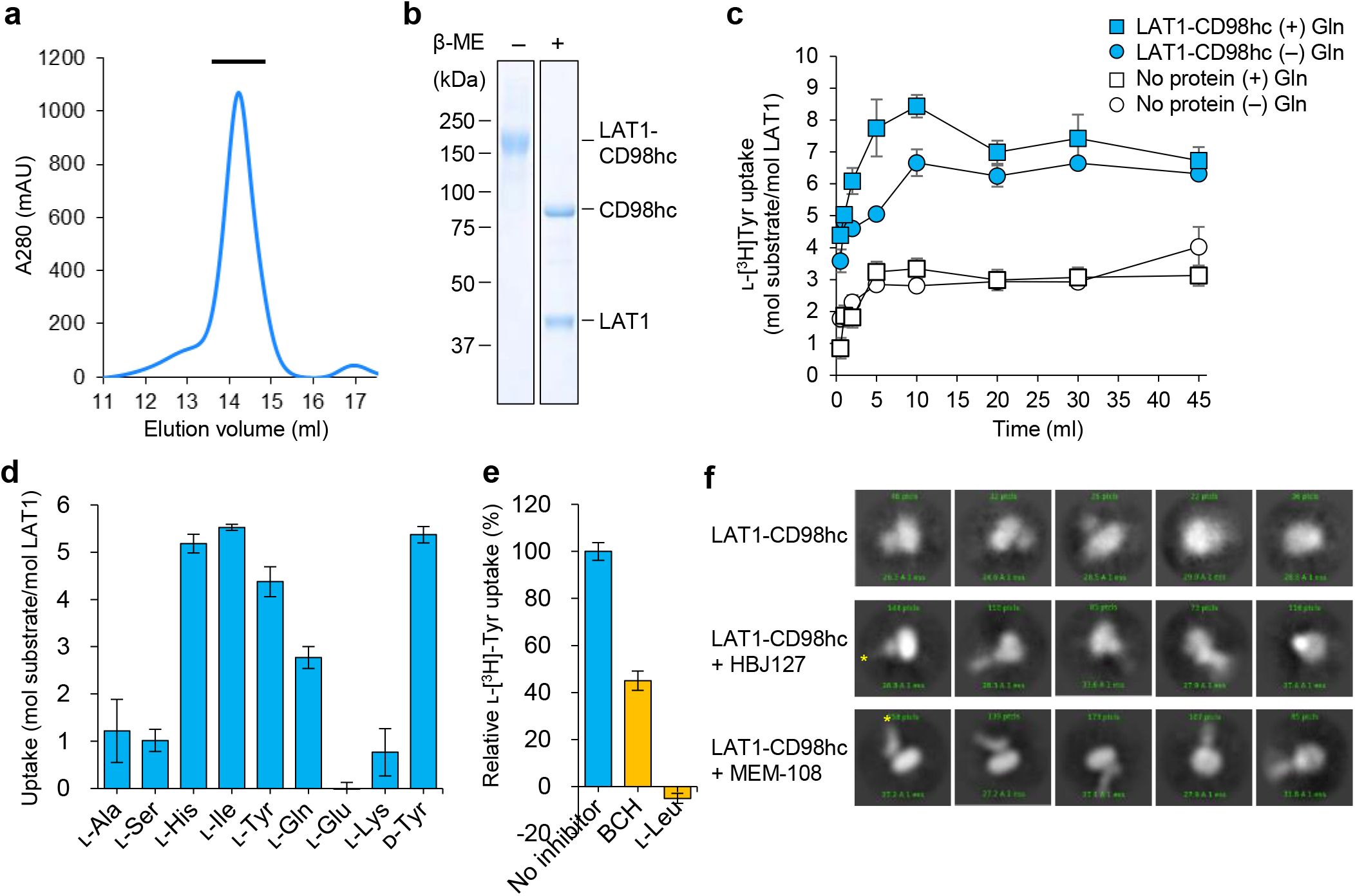
Purification and biochemical characterization of LAT1-CD98hc. a) Size-exclusion chromatography profile of LAT1-CD98hc. The fractions used for structural and functional analyses are indicated by a black bar. b) SDS-PAGE analyses of LAT1-CD98hc. Purified LAT1-CD98h was run on the SDS-PAGE gel with or without 20 mM β-ME. The band at ~160 kDa separated into the two bands at ~90 kDa and ~40 kDa, demonstrating the presence of a disulfide bond. c) Uptake of L-[^3^H]Tyr (50 μM) by purified LAT1-CD98hc reconstituted in liposomes (cyan) or by control liposomes (white). Proteoliposomes preloaded with 4 mM L-Gln (rectangle) exhibited an initial overshoot of uptake (~ 1—10 min) as compared to those without preloading (circle), indicating the antiport reaction. Values are mean ± SEM. n = 3. d) Substrate specificity of purified LAT1-CD98hc. Proteoliposomes were incubated with various radiolabeled amino acids (50 μM) and the uptake was measured at 10 min. Control liposomes were used for background subtraction. Values are mean ± SEM. n = 3. e) Inhibition assay. Proteoliposomes were incubated with L-[^3^H]Tyr (10 μM) in the presence of 30 mM competitive inhibitors in the external solution. Values are mean ± SEM. n = 3. f) Representative 2D class averages of LAT1-CD98hc with or without Fab. Cryo-EM images were recorded on a 200 kV JEM2010F microscope equipped with a CCD camera. Upper, LAT1-CD98hc; middle, LAT1-CD98hc-HBJ127 Fab; lower, LAT1-CD98hc-MEM-108 Fab. Signals corresponding to Fab molecules are highlighted by yellow asterisks. Note that the box sizes are different between the images with or without Fab.

To validate the biochemical function of the purified LAT1-CD98hc, we measured its transport activity in proteoliposomes. LAT1-CD98hc reconstituted in liposomes catalyzed the uptake of radiolabeled □-Tyr (Fig. 1c), in agreement with previous cellular assays *(2, 10).* This □-Tyr uptake exhibited an overshoot when supplemented with intra-liposomal □-Gln, demonstrating the antiport mode *(10, 19)* (Fig. 1c). We performed similar uptake experiments for different amino acids, which showed that LAT1-CD98hc transports His, Ile, Tyr (both □- and □-isomers) and Gln, but not Ala, Ser, Glu, or Lys or only weakly, consistent with the reported substrate specificity *(2)* (Fig. 1d). Although we were not able to measure the effective uptake of radiolabeled J-Leu due to the high background (Fig. S1e), the competitive inhibition assay unambiguously demonstrated the strong inhibition by □-Leu, supporting its transport by LAT1-CD98hc (Fig. 1e). We also examined the inhibition by known LAT1 inhibitors, BCH, SKN-103 and JPH203, which all resulted in significant decrease in the □-Tyr uptake, whereas SKN-102, a LAT1-insensitive analog of SKN-103 *(43)*, did not (Fig. 1e and Fig. S1f,g). These results confirmed that the purified LAT1-CD98hc is biochemically active and reflects its correct functional properties.

### Cryo-EM

LAT1-CD98hc is a relatively small (~125 kDa) membrane protein without symmetry, which makes it a challenging target for structure determination by cryo-EM *(44)*. To increase the particle size and add features for image alignment *(45)*, we utilized commercial anti-CD98 antibodies. A screening identified two mouse monoclonal antibodies, clones HBJ127 and MEM-108, as promising candidates (Figs. S2; also see Methods). These antibodies were raised against human cancers, and HBJ127 is known to have strong anti-tumor activity *(46).* Initial cryo-EM imaging of the Fab-bound LAT1-CD98hc complexes indicated that MEM-108 is more rigidly bound than HBJ127, suggesting its superior property as a structural marker (Fig. 1f). We recorded 3,382 micrographs of the LAT1-CD98hc-MEM-108 Fab complex on a 300 kV Titan Krios G3i microscope equipped with a Falcon III direct electron detector, with a small pixel size (0.861 Å/pix) to improve the signal-to-noise ratio. From a total of 361,967 particles, the 3D reconstruction yielded a map with the global resolution of 3.53 Å, according to the Fourier Shell Correlation (FSC) = 1.43 criterion. Subtraction of the micelle signal improved the overall resolution to 3.41 Å (Fig. S3 and Table S1)

The resulting EM map was of excellent quality, and resolved all of the protein components, including the extracellular domain (ED) of CD98hc, its single transmembrane helix (TM1’), all twelve transmembrane helices of LAT1 (TM1-TM12), and MEM-108 Fab. The map also resolved the disulfide bond, glycans and lipids (Fig. S4). The local resolution analysis indicated a better resolution at CD98hc-ED, which was around 3.2 Å, as compared to the TMs and Fab, where the resolution varied from 3.2-4.0 Å in the core to 4.0-4.7 Å in the periphery (Fig. S3f). The high quality of the map enabled us to build an atomic model of the LAT1-CD98hc complex mostly *de novo* (Fig. S4). The modeling of LAT1 was guided by the prominent features of the bulky side chains (Fig. S4a), and the resulting model is consistent with the evolutionary coupling analysis *(47)* (Fig. S5a). For CD98hc, we used the crystal structure of CD98hc-ED *(33)* as the starting model, and built TM1’ and the domain linker based on the map features. Since the sequence of MEM-108 is not publicly available, we could not model the side chains of its six complementarity determining regions (CDRs), and thus these regions were modeled with polyalanine. The final model includes 403 residues in LAT1, 451 residues in CD98hc and 432 residues in MEM-108 Fab.

### Architecture of LAT1-CD98hc

The cryo-EM structure of LAT1-CD98hc revealed the hitherto unobserved architecture of a transporter-glycoprotein hetero-complex, wherein the large extracellular domain (ED) of the glycoprotein CD98hc sits on top of the transmembrane domain (TMD) of LAT1, connected by a short linker (Fig. 2a, b). The dimensions of the heterodimeric complex are 90 Å in height and 60 Å in width. This architecture is consistent with an earlier topology prediction *(3)* and the previous low-resolution 3D reconstruction from LAT2-CD98hc *(48).* On the extracellular side, the ED of CD98hc is positioned above LAT1 with a ~20 Å shift from the center, creating a vertical interface. The orientation of the CD98hc-ED agrees with the previous crosslinking data *(48)* (Fig. S5b), wherein its Arg/Lys-rich surface *(33)* faces the extracellular loops of LAT1. The surface electrostatic potential calculation showed that the LAT1-facing surface of CD98hc is positively charged, whereas the corresponding surface of LAT1 is negatively charged, indicating the electrostatic interaction between the two subunits (Fig. S5c). Within the lipid bilayer, a single transmembrane helix of CD98hc (TM1’) is positioned adjacent to TM4 of LAT1, creating a lateral interface. We observe five flat-shaped densities at this interface, which could be best modeled as cholesterols (Fig. 2 and Fig. S4; discussed later). MEM-108 Fab binds to CD98hc-ED on the opposite side of the LAT1-CD98hc interface, extending towards the extracellular space by about 40 Å with a ~45° tilt.

**Figure 2.**
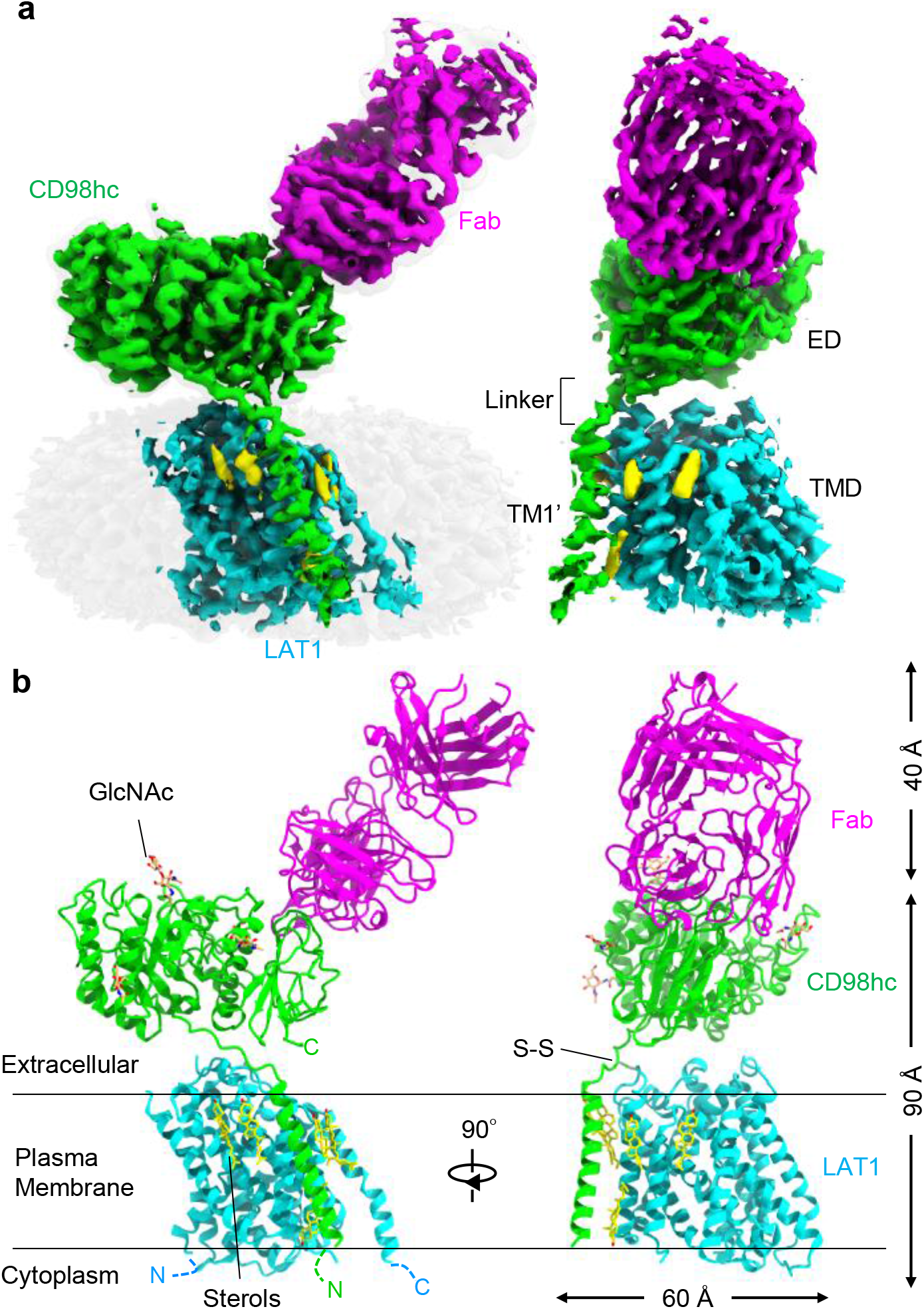
Architecture of LAT1-CD98hc. a) Cryo-EM map of LAT1-CD98hc-MEM-108 Fab. Map segments are colored based on subunit identities. The map before micelle subtraction is shown on the background. b) Atomic model of LAT1-CD98hc-MEM-108 Fab. LAT1 (cyan), CD98hc (green) and MEM-108 Fab (magenta) are shown as ribbon models. Glycans (orange), cholesterols (yellow) and disulfide bridge (green) are shown as stick models. The disordered N- and C-termini are depicted as dotted lines.

### Structure of LAT1

We first turned to LAT1, the functional transport unit of the heterodimer *(6)*. LAT1 has 12 TMs, including the ten core TMs arranged as a 5 + 5 inverted repeat topology that is typical of the APC superfamily *(5)*, plus the peripheral hairpin helices TM11 and TM12 (Fig. 3a). As in other APC members, the ten core TMs can be grouped into three functional subdomains, named hash (TMs 3, 4, 7 and 8), bundle (TMs 1, 2, 6 and 9) and arms (TMs 5 and 10) *(49).* On the extracellular side, LAT1 has six extracellular loops (EL1-6), and the intracellular side has five intracellular loops (IL1-5) and the N- and C-termini. Most of these loops are resolved in the cryo-EM map, and EL2 folds into a structured loop, EL4 forms two short helices (EL4a and EL4b) and IL1 forms one short helix (Fig. 3a, and Fig. S4 and Fig. S6).

**Figure 3.**
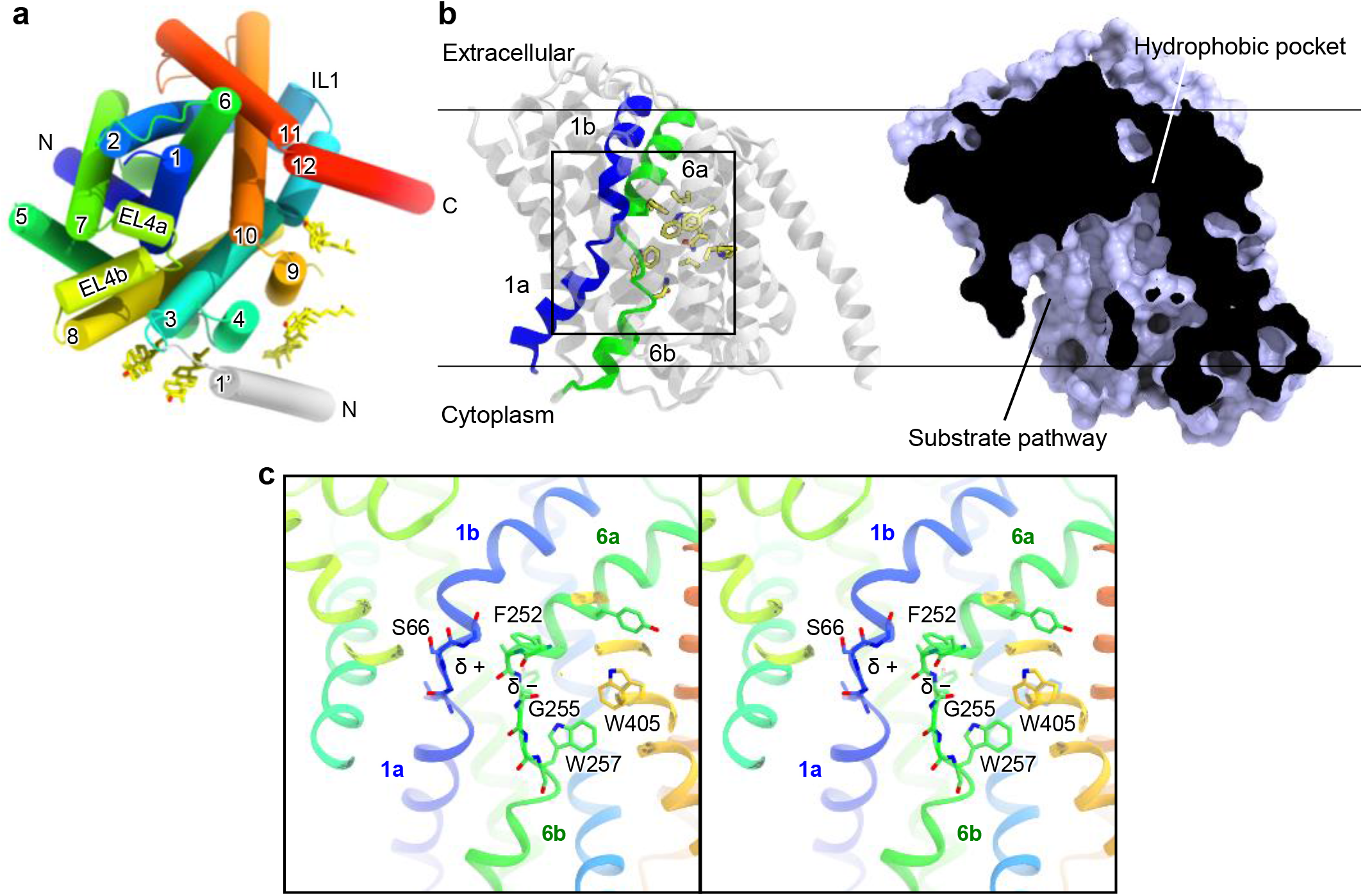
Structure of LAT1. a) Top view of LAT1 from the extracellular side. Helices are colored from blue to red from the N- to C-terminus. TM1’ of CD98hc is shown in gray. b) Side views of LAT1 in ribbon and surface representations. The helices comprising the gating bundle (TM1a, TM1b, TM6a and TM6b) are colored as in a. On the right side, cut-away surface is shown to indicate the inward-open cavity. c) Stereo view of the central substrate-binding pocket. The positive and negative poles of TM1b and TM6a are depicted with δ + and δ – symbols.

In the bundle domain, TM1 and TM6 are each divided into two discontinuous helices, TM1a and TM1b, and TM6a and TM6b (Fig. 3b). The EM map showed that the cytoplasmic portions (TM1a and TM6b) swing away towards the lipid bilayer, forming a cytoplasmic, solvent-exposed cavity (Fig. 3b). Numerous hydrophilic residues face this cavity, creating a hydrophilic substrate passageway (Fig. S6c,e). This region reportedly forms a cytoplasmic gate in the bacterial APC transporters *(49)*, and thus in the current LAT1 structure the cytoplasmic gate is open (Fig. S6). Notably, the cytoplasmic passageway is also partly open toward the lipid bilayer (Fig. 3b). A close inspection of the EM map before micelle subtraction revealed that the micelle in this region is thinner than in other regions (~28 Å), indicating a deformation of the lipid/detergent belt (Fig. S3f). This lipid-faced opening may provide a passageway for hydrophobic substrates or inhibitors.

At the end of the cavity is a conserved substrate-binding site shared by all characterized APC transporters *(50).* The structure revealed an empty space without any prominent density (Fig. 3c and Fig. S4b), indicating the apo state of the transporter. One side of this pocket is surrounded by several hydrophobic residues, such as Trp257, Trp405 and Phe252 (Fig. 3b). The extracellular side of the pocket is sealed by numerous hydrophobic and hydrophilic residues on TM1b, TM3, TM6a and TM10, which form an extracellular gate *(50)* (Fig. S6c,d). The extracellular gate is further sealed by EL4a (Fig. S6a). Above the extracellular gate is an extracellular vestibule, corresponding to an empty space between LAT1 and CD98hc (Fig. S5c). The vestibule is surrounded by numerous charged residues, which may provide a favorable environment for substrate recruitment when the extracellular gate is open. Overall, our structure captures LAT1 in the inward-open state, which features a large, hydrophobic cavity in the substrate-binding site, which is unique to this protein.

### Large, neutral substrate-binding site

An important hallmark of LAT1 is its wide substrate specificity towards bulky hydrophobic amino acids and analogs, including anti-cancer drugs such as melphalan and the thyroid hormones T_3_ and T_4_ *(20, 22).* Although our structure is in the apo state, the basic principle of substrate recognition can be deduced from a comparison with other APC superfamily transporters. To understand the broad substrate specificity, we compared the substrate-binding site of LAT1 with those of the bacterial SLC7 homologues, GkApcT *(30)* and EcAdiC *(31, 32).* These members share 25.4% and 20.3% sequence identities with LAT1, respectively, but transport smaller amino acids such as Ala and Arg. The structural comparison revealed that the TM1a-TM1b loop is almost identical between the three members, with the conserved the G(S/T)G motif in the middle (Fig. S7a,b). In ApcT and AdiC, this G(S/T)G motif creates a loop structure, which exposes the backbone amide of the second Ser/Thr to the solvent and provides a hydrogen-bond donor to the substrate carboxyl group (31). In our structure of LAT1, ^65^GSG^67^ form a similar loop structure, exposing the amide group of Ser66 towards the empty substrate-binding site (Fig. 3c and Fig. S7c). This suggests a similar manner of substrate carboxyl recognition in LAT1. In addition, the electropositive potential from the helix dipole of TM1b would provide a favorable interaction with the electronegative substrate carboxyl group (Fig. 4a, b).

**Figure 4.**
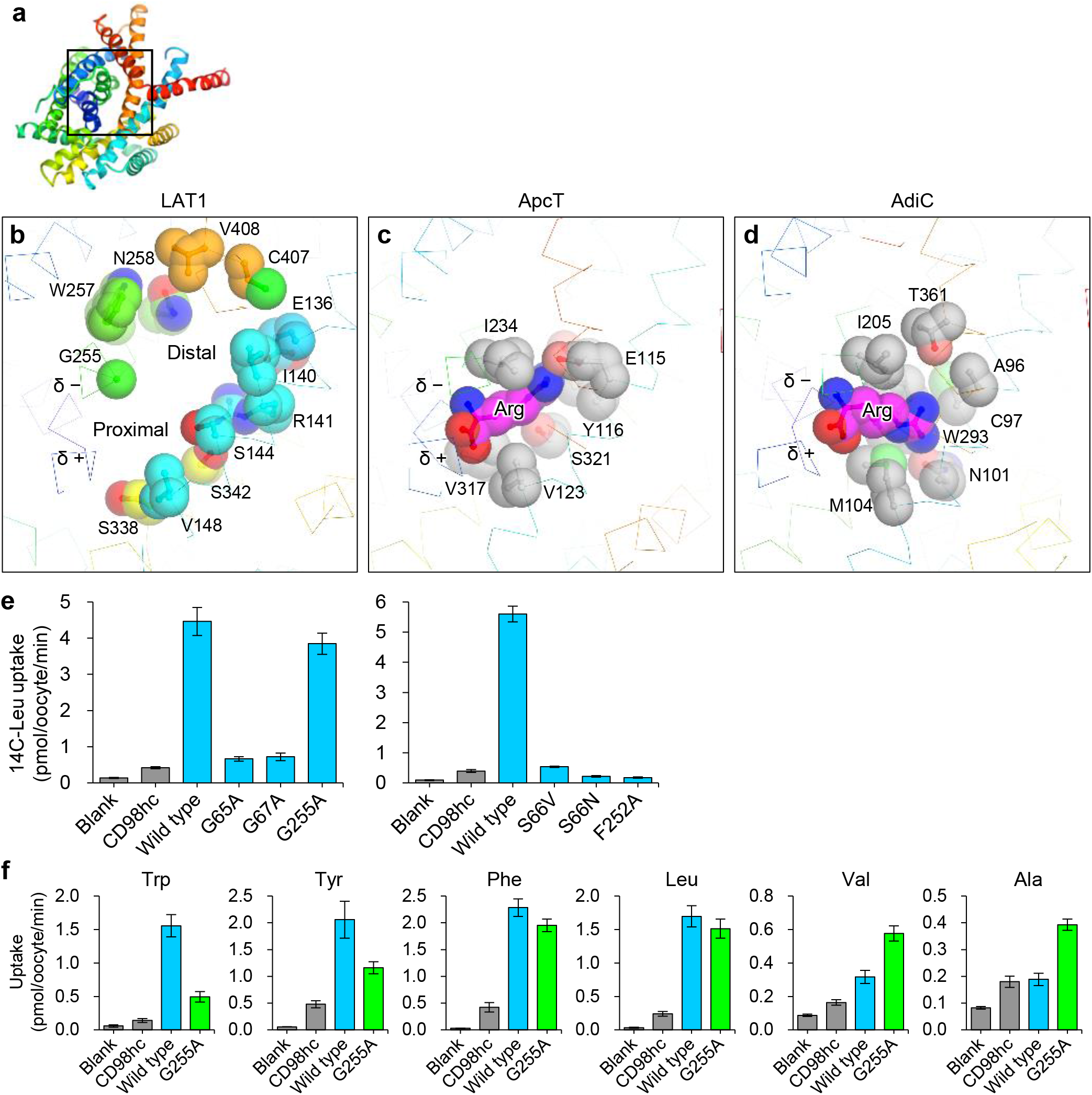
Structural basis for broad substrate specificity. a) Top view of LAT1. b) Close-up view of the substrate-binding site of LAT1. Key residues are shown as CPK models. c) Substrate-bound structure of ApcT. The Arg substrate is shown as a CPK model. Note that this structure is from the ApcT M321S mutant, which transports Arg. d) Substrate-bound structure of AdiC. e) Uptake of L-[^14^C]-Leu by *X. laevis* oocytes co-injected with the cRNAs encoding CD98hc and LAT1 variants. Blank: non-injected control, CD98hc: injected only with the CD98hc cRNA. The experiments performed on two different batches of oocytes are shown separately. Values are mean ± SEM. n = 5-9. f) Uptake of various amino acids by G255A. Values are mean ± SEM. n = 6-9.

In contrast to the TM1a-TM1b loop, the TM6a-TM6b loop is only partly conserved (Fig. S7). The loop starts with the conserved Phe/Trp motif, and its main chain carbonyl provides a hydrogen-bond acceptor for the substrate amide group in ApcT and AdiC *(30, 31)* (FigS7). In addition, the aromatic group of Phe/Trp forms a hydrophobic interaction with the substrate Cβ atom. This structure is conserved in LAT1, and thus can provide similar backbone amide and Cβ recognition sites (Fig. 3c). However, three residues after Phe252, there is Gly255 in LAT1 instead of Ile in ApcT and AdiC, which would form a sidewall to restrict the conformational freedom of the substrate side chain (Fig. 4c, d). Consequently, in LAT1 the TM6a-TM6b loop falls apart from TM10 and creates an additional pocket (named the distal pocket; Fig. 4b and Fig. S7). The distal pocket is ~17 Å long and ~14 Å wide and surrounded by a cluster of hydrophobic residues (Leu251, Trp257, Trp405 and Val408), and thus is suitable for accommodating bulky hydrophobic side chains (Fig. 4b). Interestingly, this pocket is offset by ~60° from the direction of the substrate Arg side chain in ApcT or AdiC (Fig. 4b–d). Thus, the substrates must be kinked to fit into the whole pocket, which may explain why LAT1 favors *meta* substitutions on the primary phenyl ring of the substrates *(51, 52).*

To examine the functional importance of the key residues for substrate binding, we generated single-point mutants of these residues and measured their Leu transport activities upon expression in *Xenopus laevis* oocytes *(10)* (Fig. 4e and Fig. S8).

Mutations in the GSG motif (G65A, S66V, S66N and G67A on TM1) strongly reduced the Leu uptake, consistent with their suggested role in substrate backbone recognition (Fig. 4e). F252A (TM6) also showed reduced uptake (Fig. 4e), probably reflecting the loss of the interaction between its aromatic group and the substrate Cβ atom (Fig. 3c). In contrast, the G255A mutation, which adds a methyl group near the proximal pocket (Fig. 4b), had minimal effect on the Leu transport (Fig. 4e). We reasoned that this substitution is tolerable for small amino acids. To further characterize G255A, we measured the transport activities for various amino acids, and found the reduced transport for larger amino acids, Trp and Tyr, whereas the transport of Leu and Phe was not severely affected, supporting a role of Gly255 in the recognition of bulky side chains (Fig. 4f). Interestingly, G255A showed increased transport for Ala and Val, which were not readily transported by the wild-type (Fig. 4f), suggesting that the added methyl group of G255A enabled favorable interaction with the small methyl group(s) of Val and Ala. Together, these results confirm the critical roles of the GSG motif and Phe252 in the substrate transport and the role of Gly255 in biasing the substrate specificity towards large amino acids.

### Structure of glycosylated, full-length CD98hc

The structure of full-length CD98hc revealed the previously uncharacterized features of this glycoprotein *(33).* Full-length CD98hc is composed of four regions, the N-terminal cytoplasmic tail, the single transmembrane domain (TM1’), the domain linker and the extracellular domain (ED) (Fig. 5a and Fig. S9). This arrangement is in agreement with an earlier topology prediction (53). The ED spans residues 215–630, and superimposes well on the previous crystal structure with a root mean square deviation (r.m.s.d) of 1.31 Å for 414 Cα atoms (note the different residue numbering due to the different isoforms). The ED consists of eight (β/α) repeats (domain A) and eight anti-parallel β sheets (domain C) (Fig. S9). The cryo-EM map revealed the presence of four glycans (Fig. S4), attached to Asn365′, Asn381′, Asn424′ and Asn508′ on domain A, which are not present in the previous crystal structure *(33)* (Fig. S9a, b). All of these glycosylation sites agree with the previous *N*-glycosylation predictions *(53).* Since we expressed the protein in HEK293 GnTI^−^ cells lacking a glycan processing enzyme *(54)*, the resolved sugar moieties probably represent the core acetyl glucosamine units (GlcNAc) of high mannose Λf-glycans. Notably, all of the glycans are located distal from the CD98hc-LAT1 interface, confirming that glycosylation is not directly involved in heterodimer formation *(55)* (Fig. 5a).

**Figure 5.**
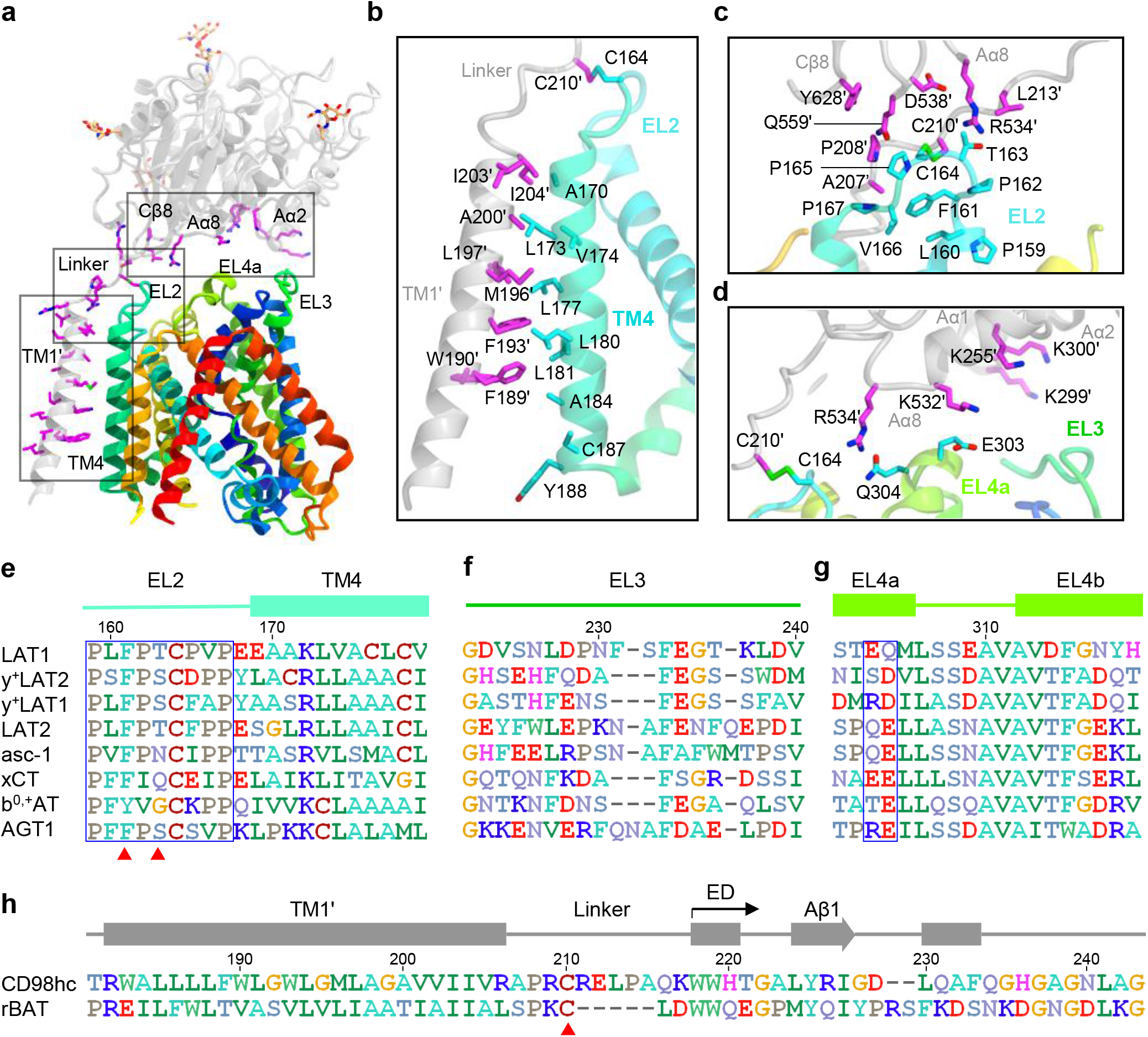
LAT1-CD98hc interface. a) The interaction between CD98hc and LAT1. b) TM1’-TM4 interface. c) Linker/Cβ2/Cβ3/Cβ8-EL2 interface. d) Aα8-EL4a and Aα1/Aα2-EL3 interface. e) Sequence alignment of EL2. f) Sequence alignment of EL3. g) Sequence alignment of EL4a-EL4b. h) Sequence alignment of CD98hc and rBAT.

The transmembrane domain of CD98hc (TM1’) spans residues 185–205, crossing the lipid bilayer with a ~10° tilt (Fig. 5a). It is positioned next to the hash motif, and forms a helix-helix contact with TM4 (Fig. 5b). We found four sterol molecules bound to this interface from the outer leaflet (Fig. S9). A previous study showed that cholesterol and its analogs increase the stability of the LAT1-CD98hc hetero-complex *(42)*. Thus, the sterol molecules we observed here explain how cholesterol stabilizes the LAT1-CD98hc interaction. In contrast to the extensive interactions in the outer leaflet, there is almost no direct contact between TM1’ and TM4 in the inner leaflet (Fig. 5b), apparently due to the mismatch of the bulky residues, Phe189′, Trp190′ and Phe193′ of CD98hc and Leu180 and Leu181 of LAT1. These bulky residues push TM1′ away from TM4 in the middle, creating a ~10 Å gap in the inner leaflet (Fig. 5b). We found one sterol density in this gap, which probably stabilizes the interface by compensating for the absence of direct interactions (Fig. S9).

The ED and TM1′ are connected by a short linker referred to as the ‘neck’ (56), which was not resolved in the previous structure of isolated CD98hc-ED *(33).* Strikingly, this linker is ordered in our structure of LAT1-CD98hc, forming an extended ~33-Å loop and bridging the ED and TM1′ (Fig. S9b). Almost all of the residues in this linker region participate in stabilizing interactions: Cys210′ forms a disulfide bridge with Cys164 of LAT1, the Arg211′ side chain forms an intramolecular salt bridge with Glu539′, the Arg211′ main chain hydrogen bonds to Asp558′ and Thr163 (LAT1), Val213′ and Pro214′ form van der Waals interaction with Trp557′ and His538′, and Arg206′ and Arg209′ interact with the interfacial sterols (Fig. S9c). Therefore, the linker of CD98hc is an integral part of the heterodimer structure rather than just a connecting peptide *(56).*

The long cytosolic N-terminal tail (~180 residues in isoform c) is not resolved in our structure, indicating a high degree of flexibility. Since the N-terminal tail has been implicated in binding with other interaction partners or ubiquitination enzymes *(29, 57)*, this flexibility may provide conformational freedom for potential interaction with these proteins. In addition, the loose packing of TM1’ to TM4 in the cytoplasmic half may allow the N-terminal tail to remain flexible even when CD98hc is associated with LAT1 (Fig. 5b).

### Heavy-light chain interface

Human CD98hc reportedly associates with six SLC7 light chains, LAT1, LAT2, y^+^LAT1, y^+^LAT2, xCT and asc-1 *(9)*. To understand the structural basis of these multi-specific heterodimerizations, we analyzed the interface between CD98hc and LAT1 in more detail and compared their sequences with those of other SLC3-SLC7 pairs (Fig. 5a–g; note that we use prime marks for the residues of CD98hc to distinguish them from those of LAT1).

There are four interfaces between the heavy chain and the light chain: TM1′-TM4, linker/Cβ2/Cβ3/Cβ8-EL2, Aα8-EL4a and Aα1/Aα2-EL3. The TM1′-TM4 interface consists of hydrophobic packing between Ile203′, Ala200′, Met196′ and Leu197′ on CD98hc and Ala170, Leu173, Val174 and Leu177 on LAT1 (Fig. 5b). These hydrophobic residues on TM4 are well conserved in the six human light chains associated with CD98hc (Fig. 5e), and probably favor similar helix-helix packing interactions. The linker/Cβ2/Cβ3/Cβ8-EL2 interface involves the disulfide bridge (Cys210′ and Cys164) and hydrogen bond (Arg511′ and Thr163), and is mediated by several direct or indirect interactions (Fig. 5c). Although it is not strictly conserved, EL2 of all light chains exhibits shared features: it is 9 residues long, it starts and ends with two conserved Pro residues (Pro159 and Pro167), Phe/Tyr is at the third position, and a hydrophilic residue is at the fifth position (Fig. 5e). These features probably enable EL2 to adopt a similar conformation in all SLC7 light chains, allowing similar interactions. In stark contrast, the CATs (SLC7A1–4), which do not associate with SLC3 *(6)*, do not share these features in EL2.

The Aα8-EL4a interface consists of two pairs of hydrophilic residues, Lys532′ and Arg534′ on CD98hc and Glu303 and Gln304 on LAT1, which participate in ionic and hydrogen-bonding contacts (Fig. 5d). Although not strictly conserved, at least one charged residue is found on EL4a of all light chains, suggesting that EL4a can form an equivalent electrostatic interaction with Aα8 (Fig. 5g). The Aα1/Aα2-EL3 interface is only partly resolved in our cryo-EM map, but there are charged residues on EL3 (Fig. 5f), which can form ionic interactions with the surface exposed residues Lys255′, Lys299′ and Lys300′ on CD98hc (Fig. 5d). In addition, Lys299′ and Lys300′ touch the surface of the micelle in the EM map before the micelle subtraction (Fig. S3f), suggesting that these charged residues may interact with the lipid head groups and function as a membrane anchor. Overall, these interfaces together contribute to the stable interaction of the heterodimer, and there is no single residue that would dictate the heterodimerization in all SLC7 members, except for the conserved Cys. This may explain the previously puzzling observation that the critical regions for heterodimer formation vary among the different SLC7 light chains *(58)*.

Notably, all of the aforementioned traits of the SLC7 light chains are also conserved in b^0,+^AT and AGT-1, which associate with another SLC3 member rBAT (SLC3A1) *(36, 59)* (Fig. 5e–f). This might imply that the association of these two light chains with rBAT is governed by different principles, such as localization or expression patterns *(37).* Meanwhile, a sequence comparison revealed that the linker of rBAT is five residues shorter than that of CD98hc, and thus the linker in rBAT cannot form an extended structure as in CD98hc (Fig. 5h). With such a short linker, the orientation of the ED of rBAT will differ from that in the LAT1-CD98hc heterodimer, and thus b^0,+^,AT- or AGT-1-rBAT may have a different interface.

### Anti-CD98 antibodies and epitopes

CD98hc is a promising target for therapeutic antibodies in cancer therapy, and HBJ127 and other antibodies indeed have antitumor activity *(60)*. During our efforts to identify suitable antibodies for the structure determination by cryo-EM, we found that HBJ127 binds to a different epitope from that of MEM-108, indicating a non-competitive binding mode. To gain further insights into the different antibody targeting mechanisms, we determined the structure of the CD98hc-ED bound to both HBJ127 Fab and MEM-108 Fab at 4.1 Å resolution (Figs S10, S11; see Methods for details). Consistent with their non-competitive binding modes, HBJ127 and MEM-108 recognize different epitopes (Fig. 6a, b). The epitope of MEM-108 involves four discontinuous sequences 410–413, 547–549, 571–575 and 617–619, which are close together in the three dimensional structure (Fig. 6c). The paratope of MEM-108 consists of CDR-H1, -H2, -H3, -L1 and -L3, which clamp the four loops of CD98hc (Fig. 6e). A local resolution analysis revealed that this antibody interface is the most ordered part of our cryo-EM map, suggesting a rigid interaction between MEM-108 and CD98hc (Fig. S11). This robust binding mode of MEM-108 suggests its potential therapeutic use.

**Figure 6.**
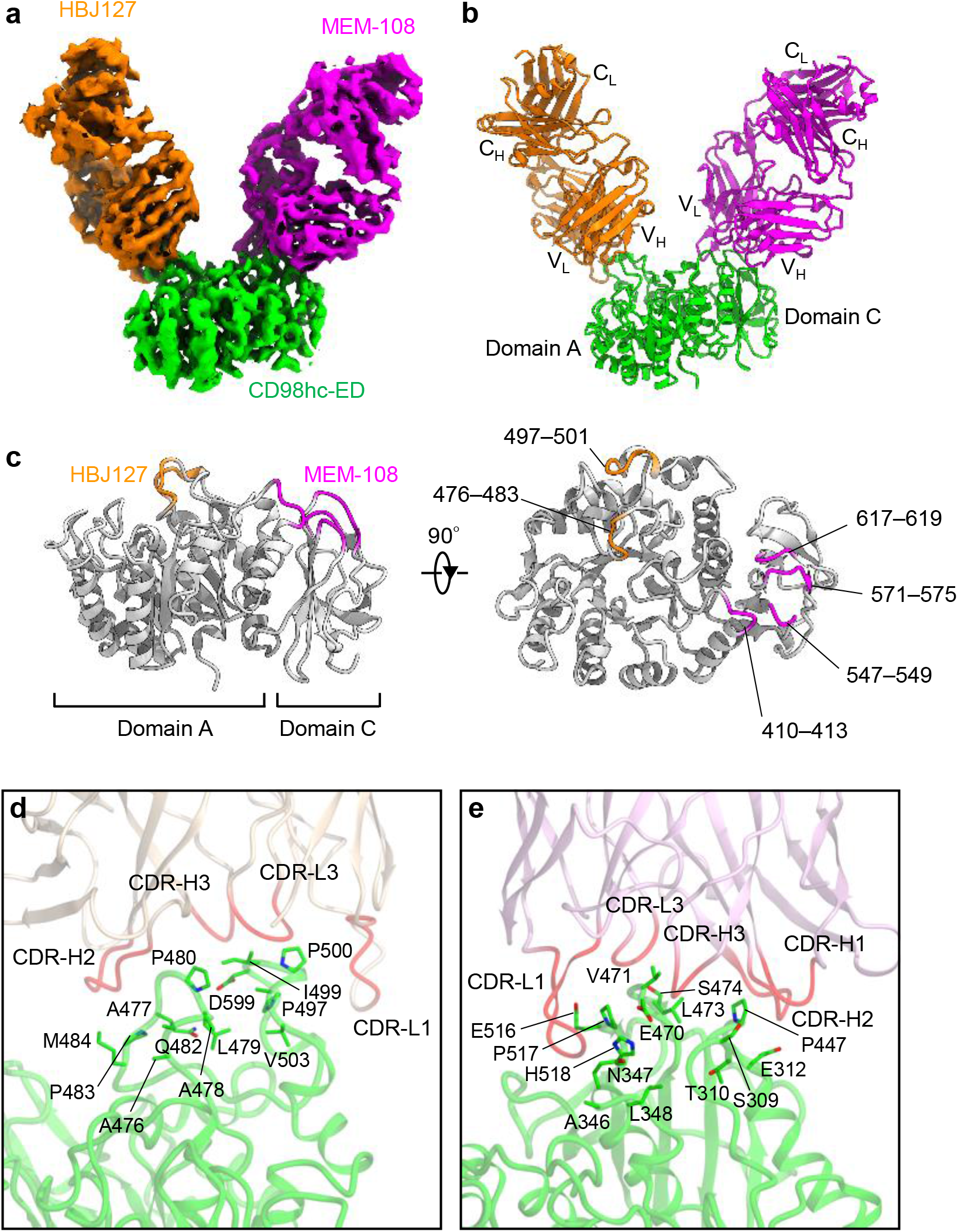
Anti-CD98 antibodies and epitopes. a) Cryo-EM map of CD98hc-ED in complex with two Fabs (HBJ127 and MEM-108). b) Structure of CD98hc-ED-HBJ127 Fab-MEM-108 Fab c) Anti-CD98hc antibody epitopes. d) CD98hc-HBJ127 interface. e) CD98hc-MEM-108 interface.

The epitope of HBJ127 involves residues 476–483 and 497–501, which are unexpectedly different from the previous data obtained from fragment-based phage-display screening *(61)* (Fig. 6c,d). Residues 476–483 are recognized by CDR-H2 and -H3, through extensive interactions involving Arg56 of HBJ127 forming ionic interactions with the backbone carbonyls of CD98hc. Residues 497–501 are recognized by CDR-L1 and -L3, involving hydrophobic interactions between Tyr31 of HBJ127 and Pro500 of CD98hc. A previous study showed that HBJ127 reacts only with human cancer cell lines and not with those from mouse or rat, indicating the absence of cross-reactivity to rodent CD98hc *(61)*. This can be rationalized by the different sequences in the second epitope loop of CD98hc in these species (residues 497–501 in human), which would disrupt the interaction with HBJ127 (Fig. S11). In addition, our structure revealed that HBJ127 only recognizes the peptide regions but not the glycans, explaining why it can robustly target various cancer cell lines that have different CD98 glycoforms *(55)*.

Together, our structure showed that these two antibodies neither disturb the structural integrity of the ED nor break the heterodimer interface. This supports the idea that most anti-CD98 antibodies act through antibody-dependent cellular cytotoxicity (ADCC) *(60)*, rather than by inhibiting amino acid transport *per se*. Nonetheless, these antibodies may still block the binding of known partners to the ED *(62, 63)*. The recently characterized humanized antibody, IGN523, exhibits robust anti-tumor activity through ADCC *(60)*, and its epitopes are predicted to be different from those presented here. Further structural studies will clarify how different antibodies exert their distinct mechanisms of action through CD98.

### Mapping SLC3/SLC7 disease mutations

The structural elucidation of the human SLC3-SLC7 heterodimer provides an opportunity to investigate the structural basis of genetic disorders associated with SLC3 or SLC7. We constructed homology models of rBAT and b^0,+^AT, in which mutations cause Type-I and non-type-I cystinuria, and y^+^LAT1, in which mutations cause lysinuric protein intolerance, and mapped the known mutations onto these structures (Fig. S12).

LAT1 has two mutations, A246V and P375L, associated with autism spectrum disorder *(39)* (Fig. S12a). A246V is located on TM6a within the extracellular gate, in a helix-helix packing interaction with TM1b. P375L is located on the cytoplasmic end of TM8, close to TM1’, TM12 and the cytoplasmic gate, or the cholesterol binding site. Thus these mutations may impair the gating process of the substrate transport. The phenylketonuria mutation G41D *(64)* exists in the unresolved N-terminal tail. Since this tail is involved in the cytoplasmic gate in other APC family members, this mutation may affect the substrate translocation process.

y^+^LAT1, or SLC7A7, has 18 missense mutations associated with lysinuric protein intolerance *(65).* Most of these mutations are clustered around the central substrate-binding site (Fig. S12b), including M50K and G54V on the TM1a-TM1b loop and S238F on the TM6a-TM6b loop. F152L is located on EL2, near the interface between CD98hc. Two other mutations, T188I and K191E, are on TM5, and probably affect the gating process of y^+^LAT1. Notably, two mutations are found on the charged residues, R333M and E251D, located on the cytoplasmic translocation pathway. These residues probably form the cytoplasmic gate according to our evolutionary coupling analysis (Fig. S5a). Therefore, mutations in either the substrate-binding site or the translocation pathway of y^+^LAT1 are pathogenic.

Mutations of rBAT, associated with type I cystinuria *(66, 67)*, are distributed across virtually the entire region of the protein (Fig. S12c). One exception is its N-terminal tail, which has no reported mutation, highlighting the importance of TM1’ and the ED for effective folding and function. Two mutations (L89P and L105R) are found within TM1’, which may impair the helix-helix interaction between b^0,+^AT and rBAT.

b^0,+^AT, or SLC7A9, has 53 known missense mutations associated with non-type I cystinuria *(66, 68).* Many mutations are found near the substrate-binding site (Fig. S12c). I44T and Y232C are located on TM1 and TM6 within the central backbone recognition site, and T123M is on TM3, in the proximity of the binding site. These mutations may severely affect substrate binding. One curious finding is that the most frequent mutations *(67)* are not located at the substrate-binding site, but are clustered around the cytoplasmic face of the transporter. For example, V170M, observed in 94% of Jewish or Israeli patients, is on IL2, and G105R, observed in 21% of white patients, is on IL1. A more pronounced mutation is P482L, which is mapped on the C-terminal tail that is disordered in our cryo-EM structure. Since these residues may form a cytoplasmic gate, further structural analyses of b^0,+^AT-rBAT are needed to understand how these frequent cystinuria mutations impair cystine transport.

## Discussion

### Agreement with previous binding-mode models

The structure of LAT1-CD98hc revealed that its substrate-binding site consists of three structural elements: (i) the positive and negative poles of the two short helices that can recognize the complementary charges of the substrate carboxyl and amide moieties, (ii) the proximal pocket that can accommodate primary side chains, and (iii) the distal pocket that provides a promiscuous binding site for hydrophobic secondary substitutions. This configuration is in striking agreement with previous hypotheses derived from pharmacophore prediction *(52)*, homology modeling (69) and structure-function analyses *(22).* We anticipate that our structure of LAT1-CD98hc at near-atomic resolution will enable more detailed investigations into the binding modes of individual substrates or inhibitors by using similar computational or experimental approaches.

### Amino acid transport

The substrate transport mechanism in the APC superfamily has been well established from more than a decade of structural and biochemical studies on prokaryotic members *(49)*. The general model is called the ‘rocking bundle’ mechanism, in which TM1a, TM1b, TM6a and TM6b undergo a series of conformational changes to transport substrates across the membrane *(70).* The structure of LAT1 confirmed that this eukaryotic APC member shares the same global architecture and functional elements as its prokaryotic counterparts, and is thus likely to follow the same rocking-bundle mechanism of amino acid transport (Fig. 7a). In the present structure, the rocking bundle is ready to accept a substrate from the cytosolic solvent (Fig. 7a). Substrate binding causes the reorientation of the helix backbones of TM1a and TM6b and eventually the major movements of the whole helices, leading to an occluded state. A possible deviation from the general model is that CD98hc-ED is on top of LAT1, and contacts EL4a. In most APC members, EL4 has been implicated in the latch function *(50)*, where it opens and closes during the transport cycle. Therefore, during the conformational changes, EL4a of LAT1 may rearrange its interaction with CD98hc, leading to a widening of the substrate pathway in an outward-open state (Fig. 7a).

**Figure 7.**
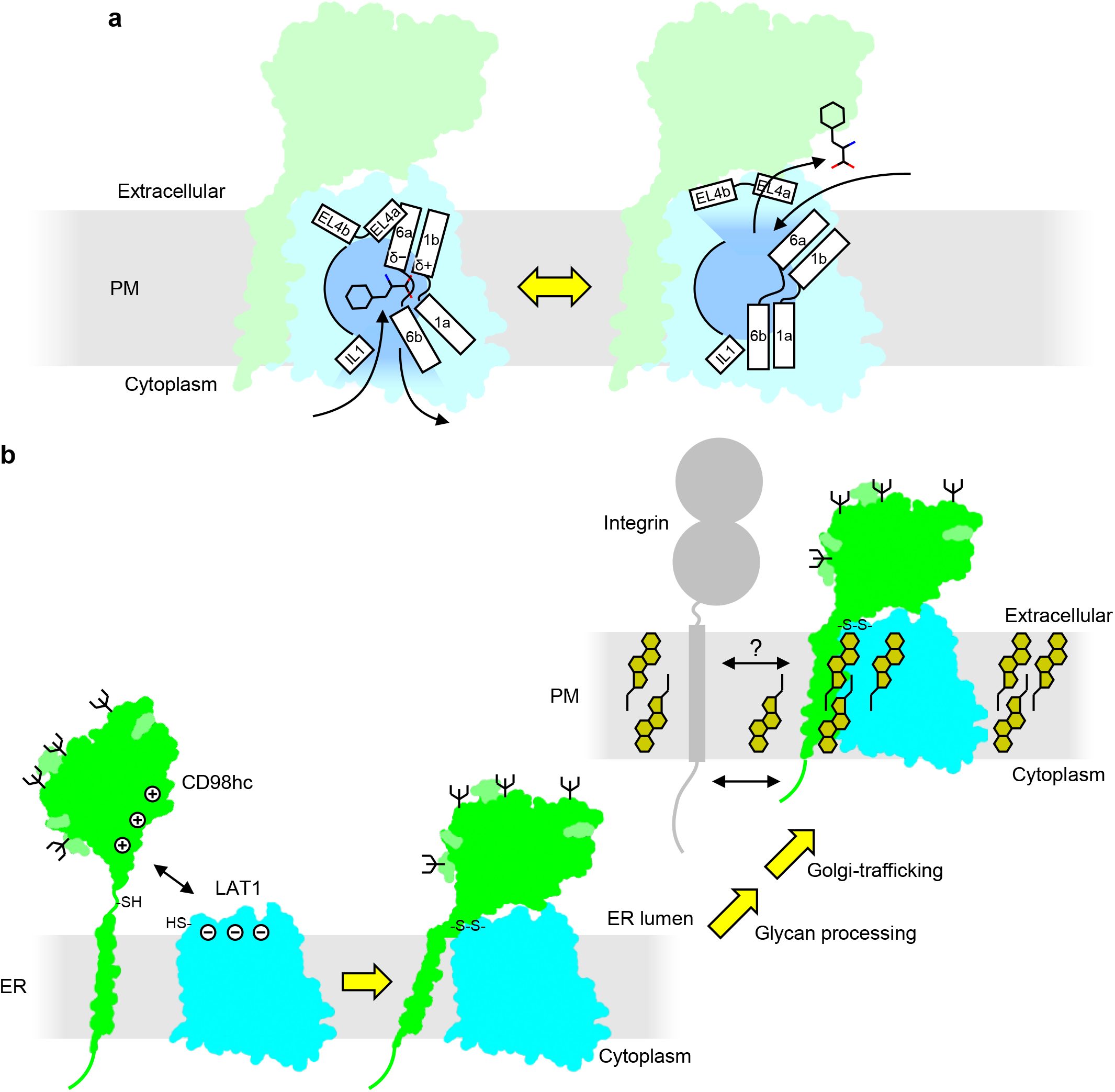
Assembly, amino acid transport and other functions of LAT1-CD98hc. a) Model of amino acid antiport by LAT1-CD98hc. b) Cartoon summarizing the biogenesis of LAT1-CD98hc. Glycosylated CD98hc assembles with LAT1 at the ER membrane, either co-translationally or post-translationally, followed by the formation of the disulfide-bond. CD98hc-ED has a positively-charged patch, which can interact with the membrane surface or LAT1. Upon assembly of LAT1, the linker of CD98hc becomes ordered. LAT1 cannot be trafficked to the plasma membrane probably due to the lack of glycosylation, and assembly with CD98hc enables the heterodimer trafficking. The heterodimer can be further stabilized upon reaching the plasma membrane by binding of cholesterols to the heterodimer interface. The integrin C-terminal tail interacts with the CD98hc N-terminal tail *(28)*, and perhaps its transmembrane domain with the CD98hc-TM1’ through the GxxxG motif.

### Integrin signaling

Apart from amino acid transport, integrin signaling through CD98hc has been well documented both *in vitro (24)* and *in vivo (25)*, and evidence supports the physical interaction between CD98hc and integrins *(29).* Previous studies have shown that although the cytosolic N-terminal tail is sufficient for the interaction, TM1’ is essential for mediating integrin signaling *(29)*. Interestingly, we found that TM1’ of CD98hc has a GxxxG motif (^195^GMLAG^199^) (Fig. S13), which is important for integrin homo- or hetero-oligomerization (71). In the present structure, this motif is not buried inside the LAT1-CD98hc interface, but is exposed to the lipid bilayer (Fig. S13). We hypothesize that this GxxxG motif of CD98hc serves as a transient or stable binding site for integrins during signal transduction *(72)* (Fig. 7b), which might explain why TM1’ of CD98hc is necessary for integrin activation.

### Assembly, trafficking and stabilization

HATs are known to undergo a complex process of assembly, maturation and trafficking *(23, 34, 73).* Building on previous biochemical data *(46)*, our structure of LAT1-CD98hc offers a near-atomic view to understand how this process proceeds (Fig. 7b). The assembly of the two chains occurs at the endoplasmic reticulum (ER) membrane, before glycan processing or Golgi trafficking, since the disulfide-liked heterodimer is detected early after translation in the core-glycosylated form *(46).* CD98hc probably folds by itself, independent of glycosylation or heterodimer formation *(33, 74)*, unlike rBAT *(34).* By using a flexible linker, CD98hc-ED can interact with either LAT1 or the lipid membrane surface *(33)* through its positively-charged patch (Fig. S5c). Upon assembly with LAT1, the flexible linker becomes fixed in a particular conformation, and the heterodimer is stabilized through multiple interactions. The stabilized heterodimer has a glycan signal on its top, which would help both LAT1 and CD98hc to exit the ER quality control *(75)* and the Golgi trafficking pathways (76). Further stabilization of the LAT1-CD98hc heterodimer would require the binding of cholesterols (Fig. S9), but the cholesterol content of the ER membrane is much lower than that of the plasma membrane (77). Therefore, the heterodimer is stabilized by being sorted into the plasma membrane, where cholesterol content can reach ~50% *(78)*. Consistent with this view, CD98 is enriched in cholesterol-rich lipid rafts *(27, 79)*.

### Other glycoprotein-solute carrier complexes

Our findings on LAT1-CD98hc may extend to other glycoprotein-solute carrier complexes *(80).* For instance, monocarboxylate transporters (MCTs; SLC16) are known to associate with CD147, another Type-II membrane glycoprotein with an immunoglobulin-like extracellular domain *(62).* Since we found that the glycoprotein CD98hc not only serves as a helper for trafficking but also as an integral part of the structure, other glycoproteins may also play structural roles in these human transporter complexes. More generally, the auxiliary glycoproteins or ‘β subunits’ have been found in various types of membrane proteins, including ion channels *(81)*, pumps *(82)* and enzymes *(83)*. However, unlike channels or enzymes that are relatively static, solute carriers must undergo rapid conformational changes during their functions. Therefore, the presence of a stable globular glycoprotein domain may affect the conformational dynamics of these solute carriers, and the extent of its influence will be an interesting subject of future investigation.

In conclusion, the cryo-EM structure of LAT1-CD98hc offers a first glimpse into the organization of a unique glycoprotein-solute carrier complex. The substrate-binding pocket of LAT1, resolved at near-atomic resolution, serves as a template for the reanalysis of the vast amount of structure-activity correlation data on LAT1 substrates and inhibitors, and will facilitate searches for better ligands through *in silico* screening. The finding that CD98hc associates with LAT1 through multiple interactions will prompt future structure-based experiments to examine which interactions play major roles in other SLC3-SLC7 pairs. An unanswered question is how the other partner proteins, such as integrins, interact with CD98hc, either as monomers or heterodimers.

## Methods

### Expression and purification of the LAT1-CD98hc heterodimer

The sequences encoding full-length human LAT1 *(SLC7A5;* UniProt ID Q01650) and CD98hc isoform c *(SLC3A2* isoform 1; UniProt ID P08195-1) were amplified from human universal reference cDNA (ZYAGEN) and cloned individually into the pEG BacMam vector *(84)*. LAT1 was fused with the C-terminal FLAG epitope, and CD98hc was fused with the N-terminal His_8_ tag and enhanced green fluorescent protein (EGFP), followed by the tobacco etch mosaic virus (TEV) protease cleavage site. Baculoviruses were generated in *Spodoptera frugiperda* Sf9 cells using the Bac-to-Bac system (Invitrogen). For expression, HEK293S GnTI^−^ cells were cultured in suspension in FreeStyle 293 medium (Gibco) supplemented with 2% fetal bovine serum. The two separate P2 baculoviruses were added at a ratio of 1:1 (v/v) at a cell density of 3–4 × 10^6^ cells/ml. The total amount of baculovirus added was ~10% (v/v) of the HEK293S GnTI^−^ culture. To boost overexpression, sodium butyrate was added at a final concentration of 5–10 mM. Cells were cultured at 37°C for 12 h after infection, and then at 30°C for another 48 h. Cells were harvested by centrifugation at 5,000 g for 12 min, resuspended in lysis buffer (50 mM Tris-HCl, pH 8.0, 150 mM NaCl, and protease inhibitors) and disrupted by probe sonication for 5 min. The lysate was subjected to ultracentrifugation at 138,000 g for 70 min, and the membrane pellet was resuspended in lysis buffer, homogenized in a glass homogenizer, and stored at −80°C.

All purification procedures were performed at 4°C. The membrane fraction was solubilized in solubilization buffer (20 mM Tris-HCl, pH 8.0, 150 mM NaCl, 1.5% dodecyl maltoside (DDM), 0.3% cholesteryl hemisuccinate (CHS), and protease inhibitors) for 90 min with gentle stirring. After removing the insoluble material by ultracentrifugation at 138,000 g for 32 min., the supernatant was incubated with ANTI-FLAG M2 Affinity Agarose Gel (Sigma) for 90 min. The gel was poured into an open column and washed with ten column volumes of wash buffer (20 mM Tris-HCl, pH 8.0, 150 mM NaCl, 0.05% DDM, and 0.01% CHS). The detergent was replaced with digitonin by washing the gel with three column volumes of digitonin buffer (20 mM Tris-HCl, pH 8.0, 150 mM NaCl, and 0.1% digitonin) and the protein was eluted by adding 100 μg/ml Flag peptide. The eluted fraction contained mostly LAT1-CD98hc but also contained free LAT1 and other unidentified proteins. To achieve more homogenous preparation, we performed another round of purification by using a GFP-nanotrap *(85)*. The GFP-nanotrap affinity resin was prepared as described previously *(86).* Briefly, an anti-GFP nanobody known as enhancer *(86)* was expressed in the *Escherichia coli* BL21 strain with an N-terminal PelB signal and a C-terminal His8 tag and purified by immobilized metal affinity chromatography. The purified nanobody was then coupled to CNBr-Activated Sepharose 4 Fast Flow beads (GE Healthcare) to yield the GFP nanotrap, with a protein-to-resin ratio of ~1 mg/ml. The Flag eluate containing the LAT1-CD98hc complex (with His8-EGFP tag on CD98hc) was reacted with the GFP-nanotrap for 90 min and washed with ten column volumes of digitonin buffer. The LAT1-CD98hc complex was cleaved off from the column by digestion with TEV protease for 16 h. The eluate was concentrated with a 100 kDa molecular weight cut-off (MWCO) concentrator (Merck Millipore) to ~3 mg/ml, as calculated from the A280 assuming that the extinction coefficient of the heterodimer was 1.33. The concentrated sample was loaded onto a Superose 6 Increase 10/300 column equilibrated with digitonin buffer. The peak fractions containing the LAT1-CD98hc heterodimer were collected and concentrated to ~2 mg/ml. The purified LAT1-CD98hc was aliquoted and stored at −80 °C.

### Transport assays in proteoliposomes

The purified LAT1-CD98hc was reconstituted in liposomes at a protein to lipid ratio of 1:120, by the polystyrene beads and dialysis method (modified from ref. *(87)*). The liposomes, composed of a 5:1 ratio of L-α-phosphatidylcholine type II-S (Sigma) to brain total lipid extract (Avanti Polar Lipids), were prepared as described *(59).* The purified proteins were mixed with Triton X-100 (0.11% w/v)-destabilized liposomes which were extruded through a 100 nm-polycarbonate filter in reconstitution buffer containing 50 mM Tris-HCl (pH 7.0) and 100 mM KCl. The detergents were then removed by stepwise additions of Bio-bead SM2 resin (Bio-Rad), followed by 16-hours of dialysis in a 10,000 MWCO regenerated cellulose membrane (Spectrum) against the reconstitution buffer. The proteoliposomes were concentrated by centrifugation at 415,000 g for 1 hr and flash frozen at −80 ^o^C until use. Empty liposomes were prepared in the same way using washing buffer instead of the purified proteins. The transport assay was performed, based on the previous experiments with another heterodimeric amino acid transporter *(59)*. Briefly, the uptake reaction was initiated by a 40-fold dilution of the proteoliposomes (~4–5 μg protein/ml) into the reconstitution buffer containing radioisotope-labeled substrates (5–20 Ci/mol), at the indicated time and concentration. The reaction was terminated by the addition of ice-cold reconstitution buffer and filtered through 0.45 μm nitrocellulose filter (Millipore), followed by washing with the same buffer. The membranes were soaked in Clear-sol I (Nacalai Tesque) and the radioactivity on the membrane was monitored with a β-scintillation counter (LSC-8000, Hitachi). For the time course experiments, prior to the transport assay, the proteoliposomes or empty liposomes were preincubated with 4 mM -glutamine for 3 h on ice, precipitated by ultracentrifugation, and then resuspended in the reconstitution buffer.

### Screening of antibodies for cryo-EM

Anti-CD98 or -LAT1 monoclonal antibodies were purchased from various commercial sources. To screen for structure-recognition antibodies, the binding properties of individual antibodies to LAT1-CD98hc were compared between native and denatured conditions. For the native conditions, purified LAT1-CD98hc was mixed with the antibody in digitonin buffer, at a molar ratio of 1:1.5, and subjected to analytical size exclusion chromatography on a Superdex 200 Increase 10/300 column. Antibody binding was assessed by monitoring the peak shift from the peak of LAT1-CD98hc. Five positive candidates identified in this assay were subjected to a negative screening by using western blotting under denaturing conditions. Purified LAT1-CD98hc was mixed with SDS-PAGE sample buffer containing 10% SDS and 20% (v/v) β-mercaptoethanol, and fractionated on the SDS-PAGE gel. The proteins were blotted with each antibody and then with corresponding secondary antibody. Among the four negative candidates that gave no detectable bands, two antibodies (clones MEM-108 and HBJ127) were selected for the current study.

For cryo-EM applications, the antibodies were digested into Fab fragments by using an IgG1 Fab preparation kit (Pierce), according to the manufacturer’s protocol. The purified Fabs were concentrated to approximately 1.5 mg/ml and buffer exchanged into digitonin buffer. For complex formation, purified LAT1-CD98hc was mixed with the Fab at a molar ratio of 1:1.2 for 1h, and then loaded onto a Superdex 200 Increase 10/300 column equilibrated in digitonin buffer. Peak fractions containing the LAT1-CD98-Fab complex were concentrated to ~5.3 mg/ml with a 30 kDa MWCO concentrator and used immediately for cryo-EM grid preparation.

### Sample vitrification and cryo-EM data acquisition

The purified LAT1-CD98hc-MEM-108 Fab (3 μl at 5.3 mg/ml) was applied onto a freshly glow-discharged Quantifoil holey carbon grid (R1.2/1.3, Cu, 300 mesh), blotted for 10 seconds at 6°C in 100 % humidity, and plunge-frozen in liquid ethane by using Vitrobot Mark IV. Vitrification of the Fab-free LAT1-CD98hc, LAT1-CD98hc-HBJ127 Fab and LAT1-CD98hc-HBJ127 Fab-MEM-108 Fab was performed with similar procedures, using different optimized sample concentrations (5.0–7.0 mg/ml) and blotting times (8–12 s). Initial cryo-EM screening was performed on a 200 kV JEM-2010F microscope (JEOL) equipped with a FastScan-F114F charge coupled device (CCD) camera (TVIPS). Imaging was performed at a nominal magnification of 80,000x, resulting in a pixel size of 1.12 Å/pix. Micrographs were recorded with an exposure time of 1–2 s, with a defocus range of −1.0 to −2.0 μm. Particles were picked in RELION-2.1 and analyzed in cryoSPARC (88).

High-resolution cryo-EM imaging of the LAT1-CD98hc-MEM108 Fab complex was performed on a 300 kV Titan Krios G3i microscope (Thermo Fischer Scientific) equipped with a GIF Quantum energy filter (Gatan) and a Falcon III direct electron detector (Thermo Fischer Scientific), in the electron counting mode. Imaging was performed at a nominal magnification of 96,000x, corresponding to a calibrated pixel size of 0.861 Å/pix. Each movie was recorded for 46.5 seconds and subdivided into 60 frames. The electron flux rate was set to 0.73 e^−^/pix/s at the detector, resulting in an accumulated exposure of 46.0 e^−^ /Å^2^ at the specimen. The data were automatically acquired using the EPU software, with a defocus range of −1.5 to −2.5 μm. A total of 3,482 movies were recorded in two separate sessions.

High-resolution cryo-EM imaging of the LAT1-CD98hc-HBJ127 Fab-MEM-108 Fab complex was performed on a 200 kV Tecnai Arctica microscope (FEI) equipped with a K2 Summit direct electron detector (Gatan), in the super-resolution mode. Imaging was performed at a nominal magnification of 23,500x, corresponding to a calibrated pixel size of 1.49 Å/pix. Each movie was recorded for 15.6 seconds and subdivided into 40 frames. The electron flux rate was set to 7.14 e^−^/pix/s at the detector, resulting in an accumulated exposure of 50.2 e^−^ /Å^2^ at the specimen. The data were automatically acquired using the SerialEM software *(89)* with a defocus range of −0.5 to −1.5 μm.

### Image processing

For the LAT1-CD98hc-MEM-108 Fab dataset, dose-fractionated movies were subjected to beam-induced motion correction using MotionCor2 *(90)*, and the contrast transfer function (CTF) parameters were estimated using CTFFIND4 *(91)*. Micrographs with poor estimated resolutions were eliminated at this point. Initially, 8,610 particles were picked from the first 28 micrographs by using the Laplacian-of-Gaussian picking function in RELION-3 *(92)* and were used to generate 2D models for reference-based particle picking. A total of 635,921 particles were subsequently picked from the 1,575 micrographs in the first session (dataset #1), and extracted with a down-sampling to a pixel size of 3.53 Å/pix. These particles were subjected to rounds of 2D and 3D classifications to select 186,605 good particles, which were then re-extracted with a pixel size of 1.38 Å/pix and subjected to 3D refinement using a soft mask covering the LAT1-CD98hc-Fab and micelle. The resulting 3D model and particle set were subjected to per-particle CTF refinement, beam-tilt refinement and Bayesian polishing *(93)* in RELION-3. The 1,807 micrographs from the second session (dataset #2) were processed similarly until particle polishing. A total of 361,967 polished particles from the two datasets were then joined and subjected to a 3D refinement, yielding a map with a global resolution of 3.56 Å according to the Fourier shell correlation (FSC) = 0.143 criterion *(94)*. To eliminate the adverse effects of the strong signals from the digitonin micelles, we performed a focused refinement *(95).* To do this, a mask covering only the protein region was prepared, by using the Volume Eraser function in UCSF Chimera *(96)*. A mask covering only the micelle belt was prepared similarly, and was used to subtract the density corresponding to the micelle belt from individual particles. A 3D refinement using the subtracted particles resulted in a nominal resolution of 3.38 Å. To further improve the map of the TMD, a 3D classification without alignment was performed by focusing on the TMD, resulting in one class (250,712 particles) with a better density at TMD. These particles were used for a final 3D refinement, which yielded a map with a 3.41 Å nominal resolution, with improved map interpretability at the TMD (Fig. S3). Local resolutions were estimated from two unfiltered half maps, using RELION-3.

For the LAT1-CD98hc-HBJ127 Fab-MEM-108 Fab dataset, motion correction and CTF estimation were performed using MotionCor2 and CTFFIND4, and subsequent image processing was done with RELION-2.1. Initially, the 2D references for particle picking were generated through rounds of manual/automatic particle picking and 2D classification. A total of 1,054,573 particles were subsequently picked from 1,354 micrographs, extracted with a down sampled pixel size of 3.54 Å/pix. After 2D classifications, 267,070 particles were selected and extracted with an original pixel size of 1.49 Å/pix. A 3D classification without any mask yielded a class containing 98,475 homogenous particles, which were then used for 3D refinement, yielding a 5.05 Å map. This map had good density at the soluble regions (the ED and two Fabs) but little interpretable density at the TMD. To better resolve the ED and Fabs, we applied a mask covering only the ED and two Fabs. By using this mask, the original 267,070 particles were subjected to a 3D classification, yielding two classes with homogenous orientations of the two Fabs relative to the ED. The 160,553 particles belonging to these classes were used for a 3D refinement with a mask covering the ED and Fabs, generating the final 4.1 Å map.

### Model building and validation

For the LAT1-CD98hc-MEM-108 Fab complex, the models of LAT1, TM1’ and the linker of CD98hc were built *de novo* into the EM density map in COOT *(97)*. The initial model of CD98hc-ED was obtained from the Protein Data Bank (ID: 2DH3), and manually fitted in COOT. In CD98hc-ED, four prominent densities were identified near four Asn residues, which corresponded to the four N-glycosylation sites predicted by the NetGlyc server (http://www.cbs.dtu.dk/services/NetNGlyc). One or two acetyl glucosamine (GlcNAc) units were built into these densities, by using the Carbohydrate module in COOT *(98).* Within the TMD, five non-protein densities were observed, exhibiting the characteristic shape of a flat sterol backbone. These densities may belong to digitonin, CHS or cholesterol, which were present during the expression and purification. We modelled these densities as cholesterol, since it represents a sterol backbone moiety shared by all candidates. For MEM-108 Fab, the initial model used was Fab 81-7 in the insulin receptor (PDB ID: 4ZXB). This Fab belongs to the same isotype IgG_1_ κ as MEM-108 (and also has the closest sequence to HBJ127; see below). The Fab was first fitted into the density as a rigid body, and the side chains in six CDRs were mutated to alanine in COOT. The main chain traces are then corrected, with some regions requiring amino acid insertions. The side chains outside the CDRs also showed deviation from the registered sequence, indicating the sequence differences between MEM-108 and Fab 81-7. Models were iteratively refined by using phenix.real_space_refine (99) in PHENIX *(100)* and manual refinement in COOT. The validation check for model stereochemistry was performed in MolProbity *(101)* (Table S1).

For the LAT1-CD98hc-HBJ127 Fab-MEM-108 Fab complex, the initial model was built from LAT1-CD98hc-MEM-108 Fab and fitted into the density with COOT. To model HBJ127 Fab, we first searched for the PDB entry with the closest sequence to the deduced sequence of HBJ127 *(102)*, by using NCBI BLAST. The search identified the mouse IgG_1_ κ antibody Fab 83-7 for Insulin receptor (PDB ID: 4ZXB) as the top hit. This Fab was fitted into the density, and amino acid residues in the CDRs were manually substituted and refined in COOT. Real-space refinements and model validation were performed in the same manner as for LAT1-CD98hc-MEM-108 Fab (Table S1).

The refined models were tested for potential overfitting using a cross-validation method described elsewhere *(103).* Briefly, the final models were ‘shaken’ by introducing random shifts to the atomic coordinates up to 0.5 Å, and were refined against the first half map. These shaken-refined models were used to calculate the FSC against the same first half maps (FSC_haf1_ or work), the FSC against the second half maps not used for the refinement (FSC_half2_ or free) or the FSC against the summed maps (FSC_sum_), using phenix.mtriage *(104).* The marginal differences between the FSC_work_ and FSC_free_ curves indicated no severe overfitting of the models (Fig. S3). All molecular graphics figures were prepared with CueMol (http://www.cuemol.org).

### Evolutionary coupling analysis

The evolutionary coupling analysis was performed using EVcoupling *(47)*, which predicts the residue-residue contacts by using the information about the amino acid sequence co-variation. Calculations were done on the web server (http://evfold.org/evfold-web/evfold.do), with all job configurations set to the default values. Initially, a multiple sequence alignment of LAT1 homologs was generated using the iterative hidden Markov model-based search tool jackhmmer *(105).* The optimal E-value cut-off was automatically determined to be −3, and the resulting alignment contained 85,721 sequences. After excluding the amino acid positions with more than 30% gaps in the column, residues 46–474 were used for calculating the evolutionary couplings by a pseudo likelihood maximization approach *(106).* The top ranked contact pairs are summarized in Fig. S5.

### Transport measurements using *X. laevis* oocytes

Expression plasmids for LAT1 and CD98hc were constructed previously *(107).* Mutations of LAT1 were introduced by whole plasmid PCR using PrimeSTAR MAX DNA polymerase (Takara) according to the manufacturer’s protocol. Amino acid substitutions to alanine (for G65A, G67A, F252A, G255A), valine (for S66V), and asparagine (for S66N) were performed by altering the corresponding codons into GCC, GTG, and AAC, respectively. cRNAs were synthesized *in vitro* from the linearized plasmids using an mMessage mMachine Kit, polyadenylated with a Poly(A) Tailing Kit and purified with a MEGAclear Kit (Ambion).

For *X. laevis* oocyte expression, defolliculated oocytes were co-injected with the polyadenylated cRNA of LAT1 and CD98hc (25 ng/oocyte). Control oocytes were injected with either CD98hc cRNA alone or water. Transport measurements were performed 2 days after injection as previously described *(2).* In brief, the oocytes were incubated for 30 min at room temperature with 500 μl Na^+^-free uptake buffer (96 mM Choline-Cl, 2 mM KCl, 1.8 mM CaCl_2_, 1 mM MgCl_2_, and 5 mM HEPES (pH 7.5)) containing the indicated concentrations of ^14^C-labeled leucine, alanine, tryptophan, tyrosine, or phenylalanine, or H-labeled valine. The radioactivity was determined by liquid scintillation counting. Reproducibility of the results was confirmed by independent experiments using different batches of oocytes.

### Plasma membrane localization of LAT1-CD98hc in *X. laevis* oocytes

For isolating the total membranes of *X. laevis* oocytes, 33 oocytes expressing CD98hc with or without LAT1 (wild type or mutants) were homogenized in 660 μl buffer A (20 mM Tris-HCl pH 7.4 and cOmplete protease inhibitor cocktail (Roche)). Cell debris and nuclei were removed by centrifugation at 1,000 g for 15 min, and the supernatant was carefully collected without transferring the yolk floating on the surface. This step was repeated three more times to avoid contamination by yolk in the supernatant. Next, 1M NaCl was added to solubilize the yolk slightly remaining in the supernatant, and 350 μl of each sample was used to collect total membranes by ultracentrifugaion at 150,000 g for 1 hour. The total membranes were solubilized in 63 μl buffer A containing 1 % Fos-Choline-12 (Anatrace) and incubated on ice for 30 min. The same volume of each sample was resuspended in SDS-PAGE sample buffer and analyzed by SDS-PAGE and immunoblotting. The antibodies used for the detection of LAT1 and CD98hc were as follows: anti-LAT1 antibody (1:2,000, J-Pharma), anti-CD98hc antibody (1:1,000, H-300; Santa Cruz Biotechnology), peroxidase goat anti-mouse IgG and peroxidase goat anti-rabbit IgG (1:10,000, Jackson ImmunoResearch).

For the immunofluorescence detection of LAT1 (wild type and mutants) and CD98hc expressed in *X. laevis* oocytes, paraffin sections were prepared as described previously *(108)*, with minor modifications. Briefly, 2 days after the injection of cRNAs, *X. laevis* oocytes were fixed in 4% paraformaldehyde-phosphate buffer (PFA/PB; pH 7.4) at 4°C overnight. After fixation, the oocytes were washed twice to remove the PFA and embedded in 2% agarose in phosphate buffer for sectioning. Samples were dehydrated in a graded alcohol series, embedded in paraffin, and cut into 5-μm thick sections. Staining was performed as described previously *(109).* After antigen retrieval with target retrieval solution (pH 9.0; Dako, Glostrup, Denmark), the slides were blocked using Blocking One Histo (Nacalai Tesque, Kyoto, Japan) for 20 min, followed by an incubation with the primary antibody, anti-LAT1 (1:500, J-Pharma), overnight at 4°C. After washing with TBS, the slides were incubated with Alexa Fluor 488-conjugated anti-mouse IgG (A21202, 1:1000, Invitrogen) for 1 hour at room temperature. Sections were then washed in TBS and mounted. Images were acquired using a fluorescence microscope (BZ-9000, Keyence) equipped with a × 100 objective lens (CFI Plan Apo λ, NA 1.40, Nikon).

### Homology model generation and mutation mapping

Homology models were generated in Modeller *(110)* by using the multiple sequence alignment of the eight human SLC7 members or that of CD98hc and rBAT. Missense mutations were summarized from the Human Gene Mutation Database (http://www.hgmd.cf.ac.uk) and the published data *(39, 64–68)*

### Data availability

The raw micrographs of LAT1 -CD98hc-MEM 108 Fab and LAT1-CD98hc-HBJ127 Fab-MEM-108 Fab have been deposited in the Electron Microscopy Public Image Archive under accession numbers EMPIAR-XXXXX and EMPIAR-XXXXX. The cryo-EM density maps have been deposited in the Electron Microscopy Data Bank under accession numbers EMD-XXXX and EMD-XXXX. Atomic coordinates have been deposited in the Protein Data Bank under accession numbers XXXX and XXXX.

### Author contributions

Y.L. and O.N. conceived the study. Y.L. performed expression screening, purified LAT1-CD98hc and Fab, prepared the cryo-EM samples and determined the structure. Y.L., T.K., T.Ni., R.I., T.Y., and M.S. collected the cryo-EM data. P.W. and S.N. performed the transport assays using proteoliposomes. C.J., L.Q., R.O., S.O. and Y.K. performed the transport assays using *X. laevis* oocytes. T.Na assisted with high-resolution cryo-EM data processing with RELION. K.O. purified the proteins for biochemical assays. H.E. provided LAT1 inhibitors and assisted with data interpretation. Y.L. and O.N. wrote the manuscript with contribution from all authors. O.N. supervised the project.

## Acknowledgements

We thank R.Danev and M.Kikkawa for setting up the cryo-EM infrastructure; K.Ogomori and M.Miyazaki for technical assistance; G.Kasuya, M.Fukuda and R.Taniguchi for discussions; and W.Kühlbrandt for comments on the manuscript; Y.L. was supported by the Toyobo Biotechnology Foundation Fellowship. This work was supported in part by MEXT/JSPS KAKENHI under Grant Numbers JP16J07405 to Y.L. and JP16H06294 to O.N.; by AMED under Grant Numbers JP18am0101082 to M.S., JP18gm0810010 to S.N., JP18cm0106131 to Y.K. and JP18am0101115 to O.N.

## Declaration of interest

The authors declare no competing interest.

