## Supplementary Figures for "Cryo-EM structure of the human L-type amino acid transporter 1 in complex with glycoprotein CD98hc"

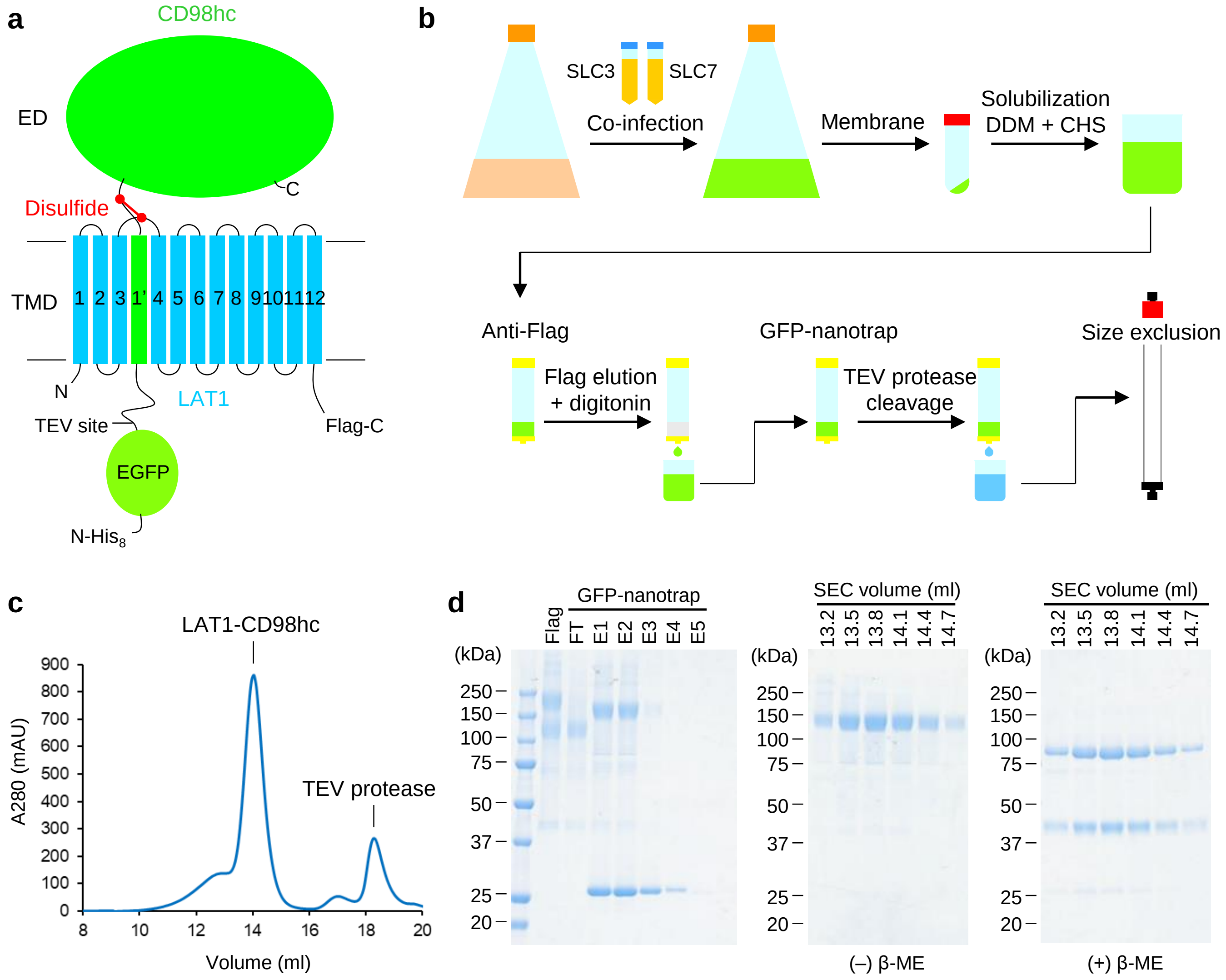

**Figure S1 | Expression, purification and biochemical assay of LAT1-CD98hc.**

a) The domain organization of LAT1-CD98hc and construct design.
b) Expression and purification strategy.
c) Representative size-exclusion chromatography profile of LAT1-CD98hc.
d) SDSPAGE analyses of LAT1-CD98hc at different stages of purification.

(Continues on the next page)

Lee et al. Figure S1

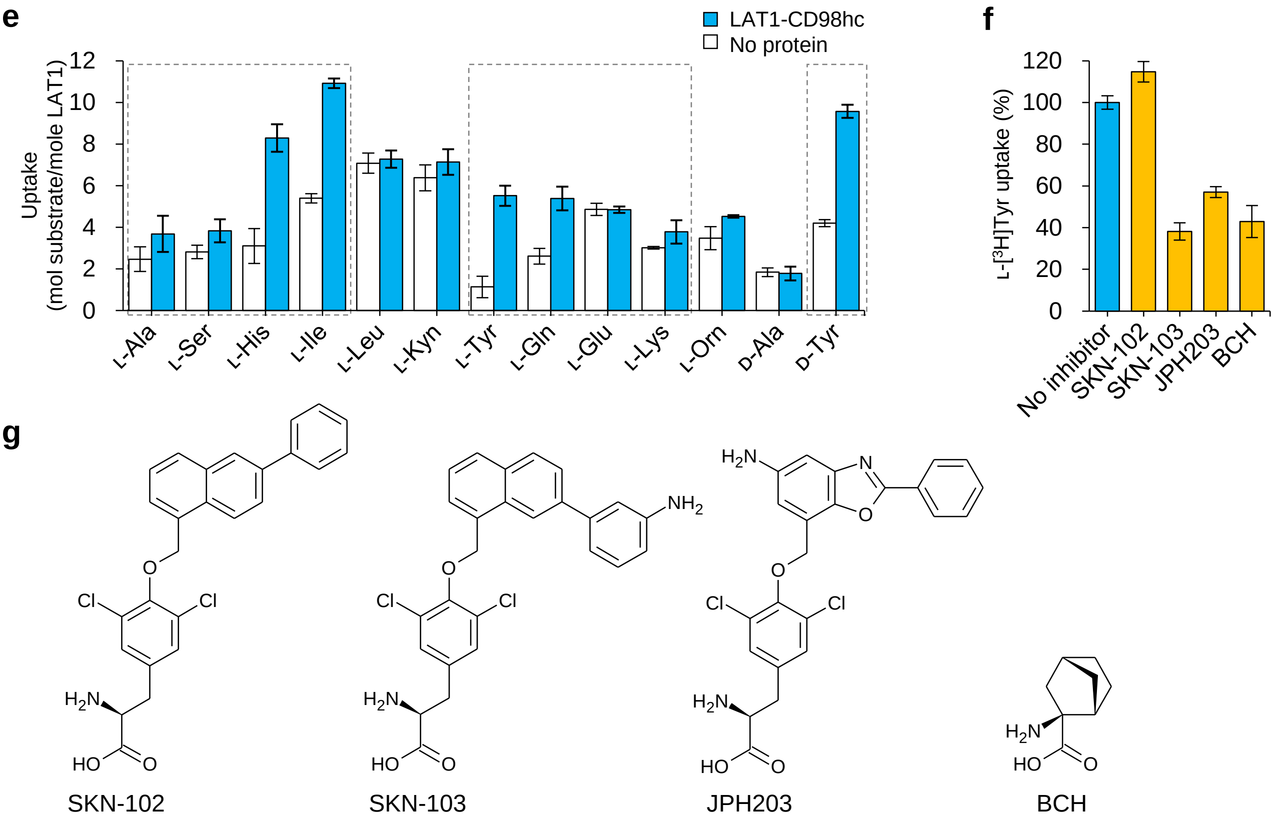

**Figure S1 | continued**

e) Substrate specificity of LAT1-CD98hc. Uptake of $^{3}$H-labeled amino acids (50 μM) was measured for the proteoliposomes reconstituted with LAT1-CD98hc or for control liposomes. The data used in Fig. 1d are marked by gray dotted rectangles.

f) The transport inhibition by LAT1 inhibitors. LAT1-CD98hc-reconstituted proteoliposomes were pre-incubated with the indicated compounds and the uptake of L-[$^{3}$H]Tyr (100 μM) was measured. The inhibitor concentrations were 30 μM for SKN-102, SKN-103 and JPH203 and 30 mM for BCH.

g) The chemical structures of LAT1 inhibitors and analogs.

Lee et al. Figure S1 (continued)

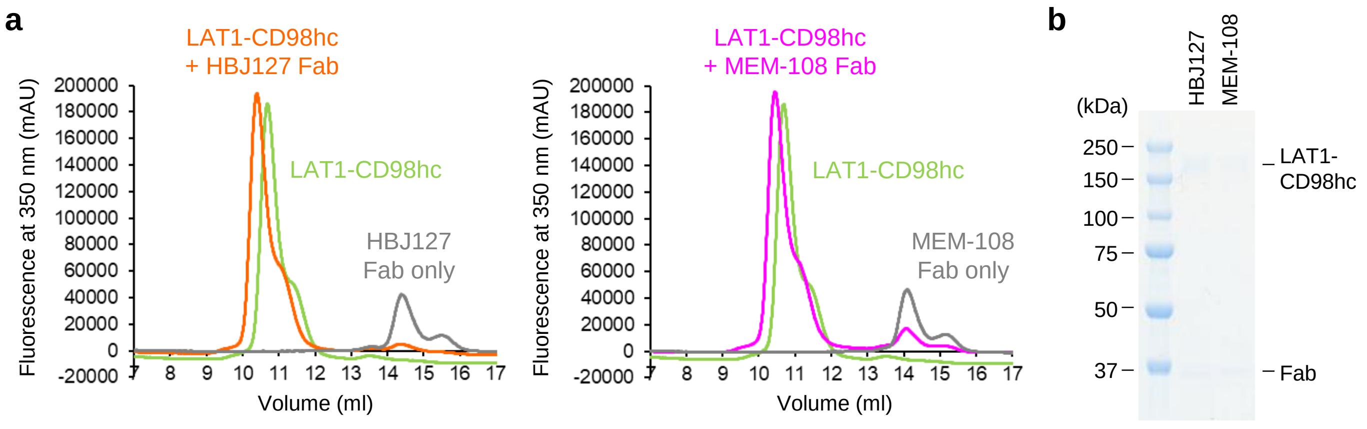

**Figure S2 | Fab complex formation of LAT1-CD98hc.**

a) FSEC analysis of the LAT1-CD98hc-Fab complexes. The traces are tryptophan fluorescence (excitation, 280 nm; emission, 350 nm).

b) SDS-PAGE analysis of the LAT1-CD98hc-Fab complexes.

Lee et al. Figure S2

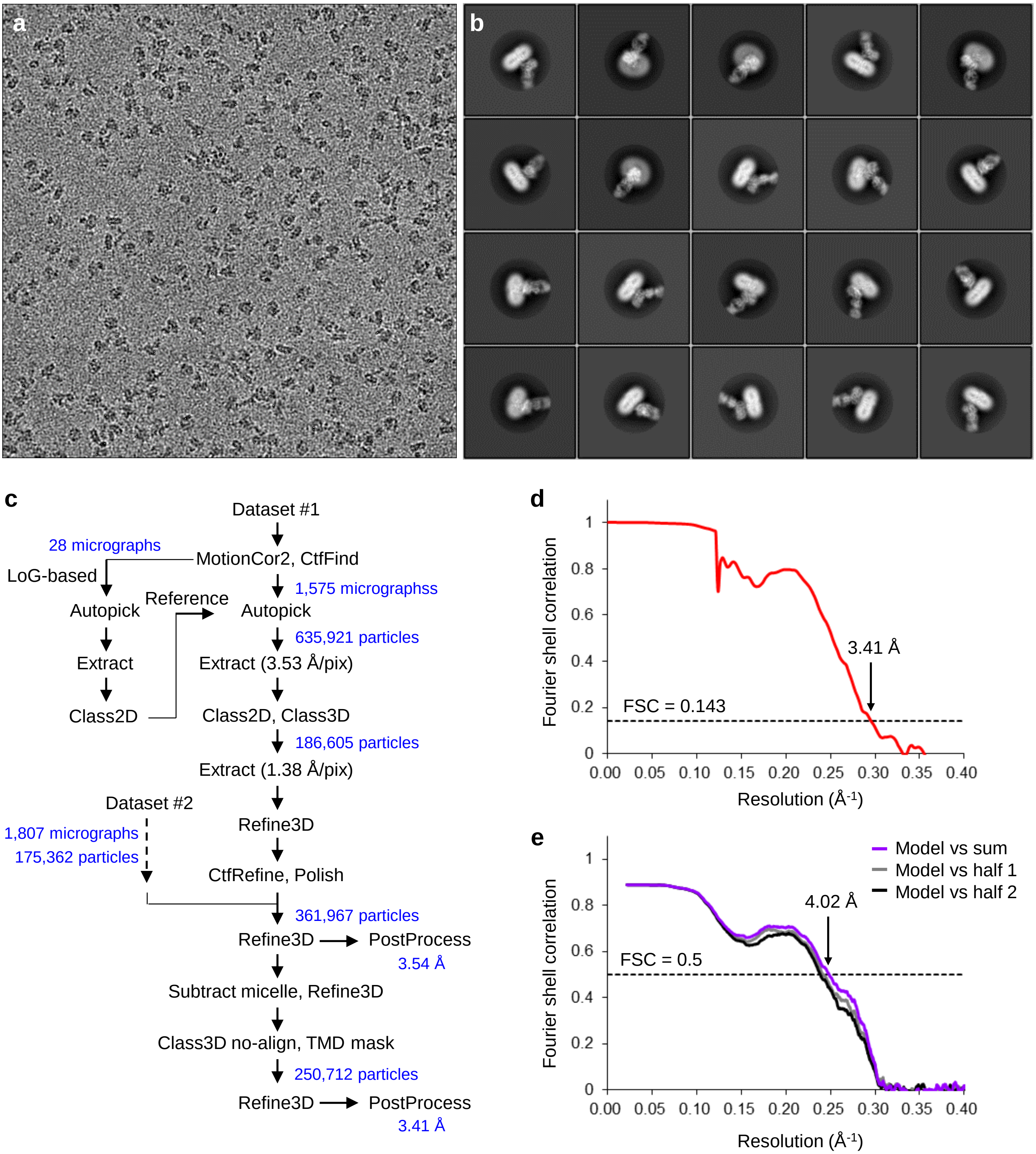

**Figure S3 | High-resolution cryo-EM of LAT1-CD98hc-MEM-108 Fab**

a) Representative micrograph of LAT1-CD98hc-MEM-108 Fab, recorded on a 300 kV Titan Krios microscope equipped with a Falcon III camera.

b) Representative 2D class averages.

c) Workflow of single-particle image processing.

d) Fourier shell correlation curve of two half-maps.

e) Map-to-model correlation.

(Continues on the next page)

### Lee et al. Figure S3

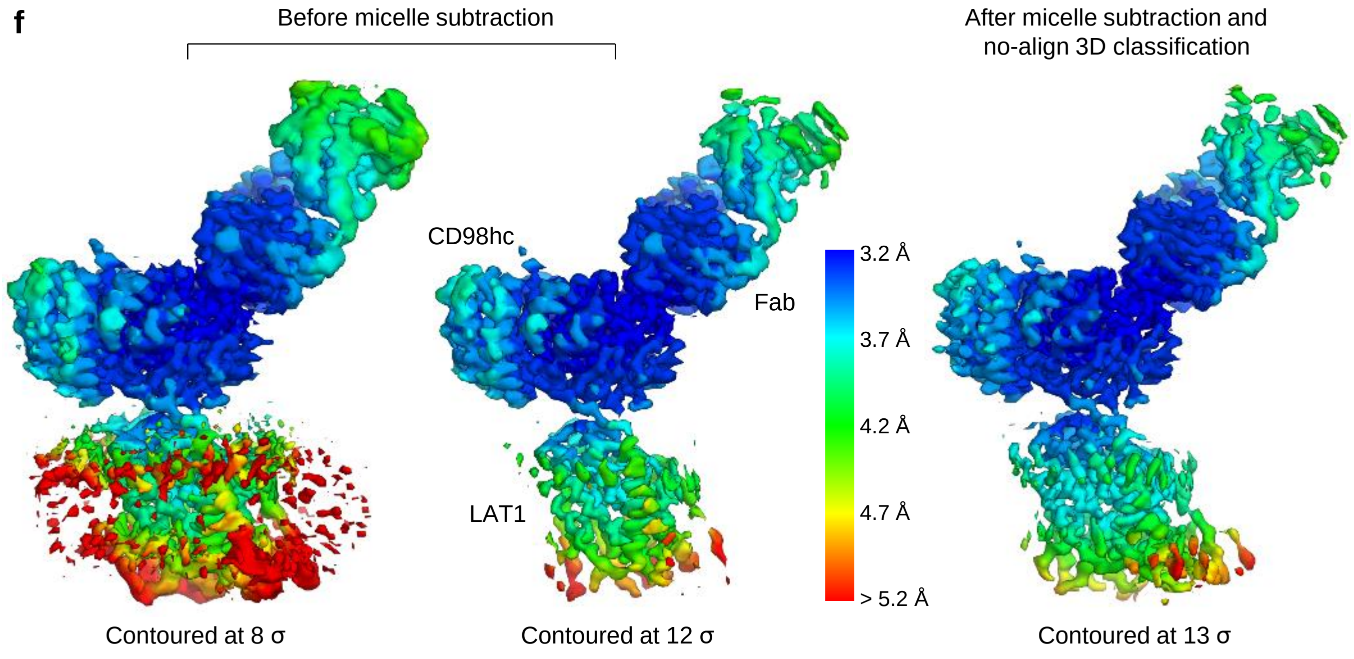

**Figure S3 | continued**

f) Local resolution analysis. The EM maps before micelle subtraction (left and center) and after micelle subtraction and no-align 3D classification (right) are shown at different contour levels. Local resolutions were calculated in RELION and colored from blue (3.2 Å) to red (5.2 Å).

Lee et al. Figure S3 (continued)

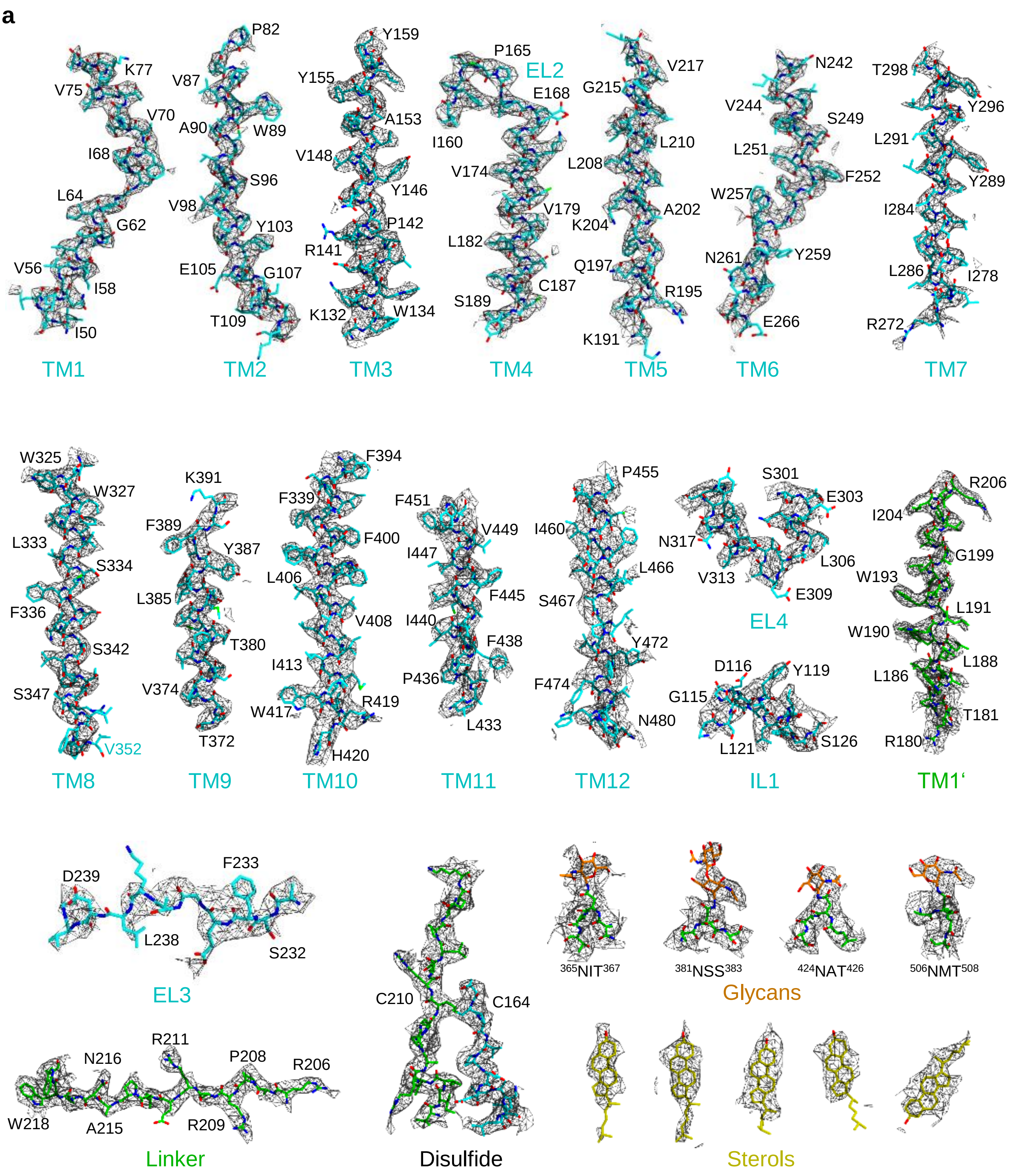

**Figure S4 | Atomic model of LAT1-CD98hc in the cryo-EM density map.**

a) Cryo-EM density maps and atomic models are shown for selected regions. TM1–TM12, IL1 and EL2–EL4 of LAT1 (cyan), TM1’ and the linker of CD98hc (green). The disulfide bond, four *N*-glycans (orange) and five lipids (yellow).

(Continues on the next page)

Lee et al. Figure S4

b

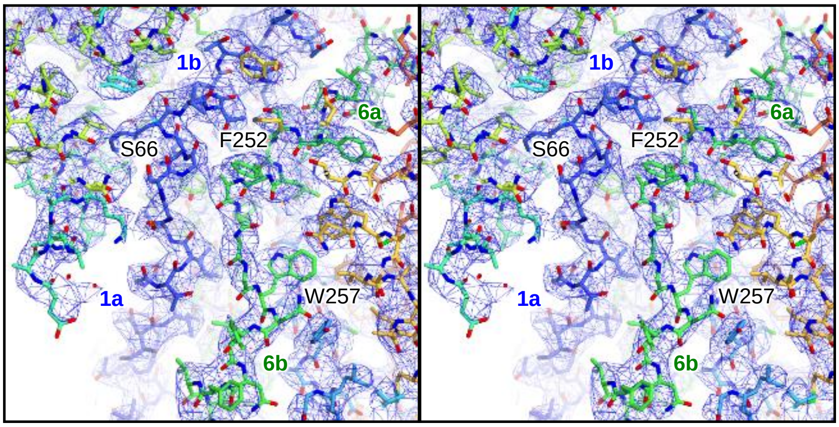

**Figure S4 | continued**

b) Stereo view of the central pocket of LAT1. The EM map shows no prominent density in the conserved substrate-binding site, indicating the apo state. The view is the same as in Fig. 3c.

Lee et al. Figure S4

**a**

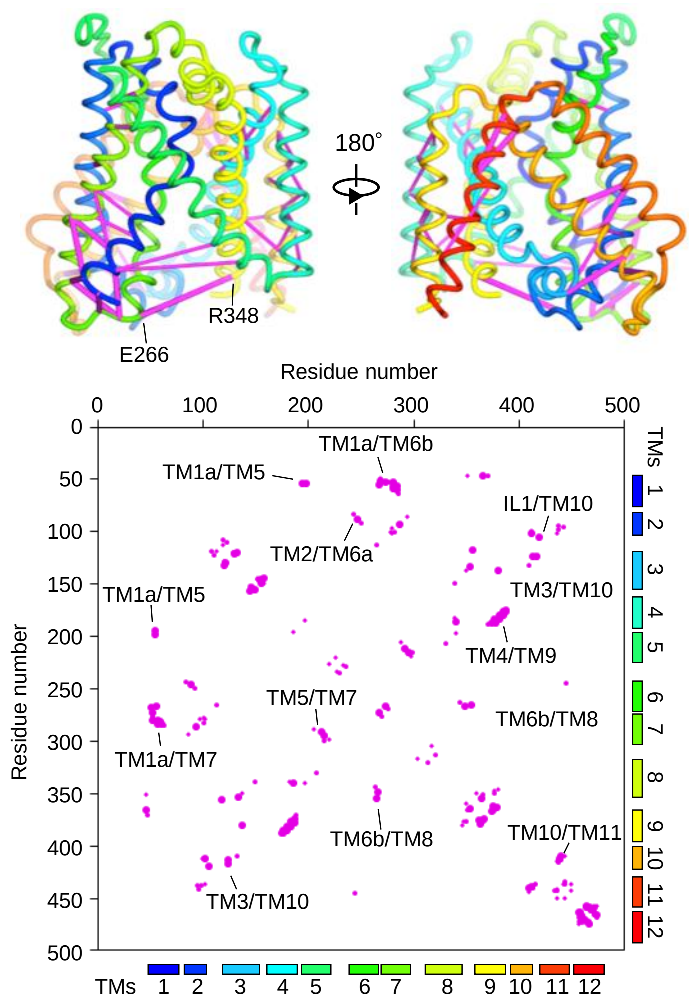

**b**

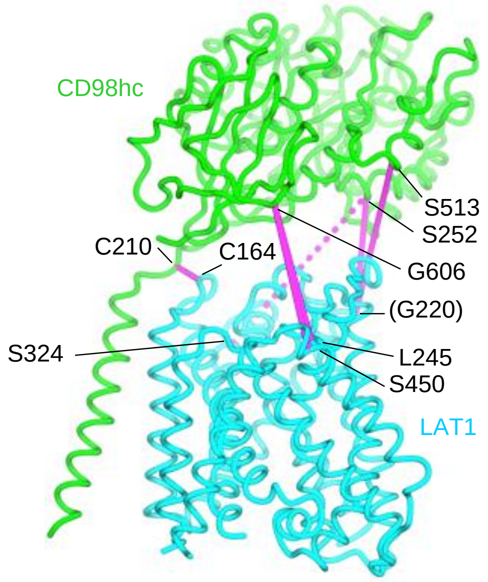

**Cross-linked pairs (ref. 48)**

| Cross-linking | CD98hc | LAT2 | LAT1 | Cα-Cα |
|---|---|---|---|---|
| Native | C210 | C154 | C164 | 5.7 Å |
| 80%-95% | S252 | C210 | G220 | N/A |
| | S513 | C210 | G220 | N/A |
| | G606 | A235 | L245 | 24.0 Å |
| | G606 | S441 | S450 | 25.5 Å |
| 60% | S252 | A315 | S324 | 34.6 Å |

**Figure S5 | Evolutionary coupling, cross-linking and electrostatic potential of LAT1-CD98hc.**

a) Evolutionary coupling analysis of LAT1. Upper, the top 50 predicted contact pairs are shown on the structure as lines. Thicker lines indicate stronger coupling. E266 and R348, which are ~19Å apart in the structure, show strong evolutionary coupling, suggesting that they contact in different conformations (*47*). Lower, the top 150 pairs are summarized in a map. Selected helix-helix contacts are indicated with labels.

b) Correlation with previous cross-linking data on LAT2-CD98hc (*48*). Upper, cross-linked pairs are shown on the structure as lines. Dotted lines are for a 60% cross-linked pair. Lower, cross-linked pairs in CD98hc and LAT2 are summarized in a table, along with the corresponding residues in LAT1 and Cα-Cα distances. N/A, not applicable because G220 is disordered, and for visualization I219 is used instead.

(Continues on the next page)

Lee et al. Figure S5

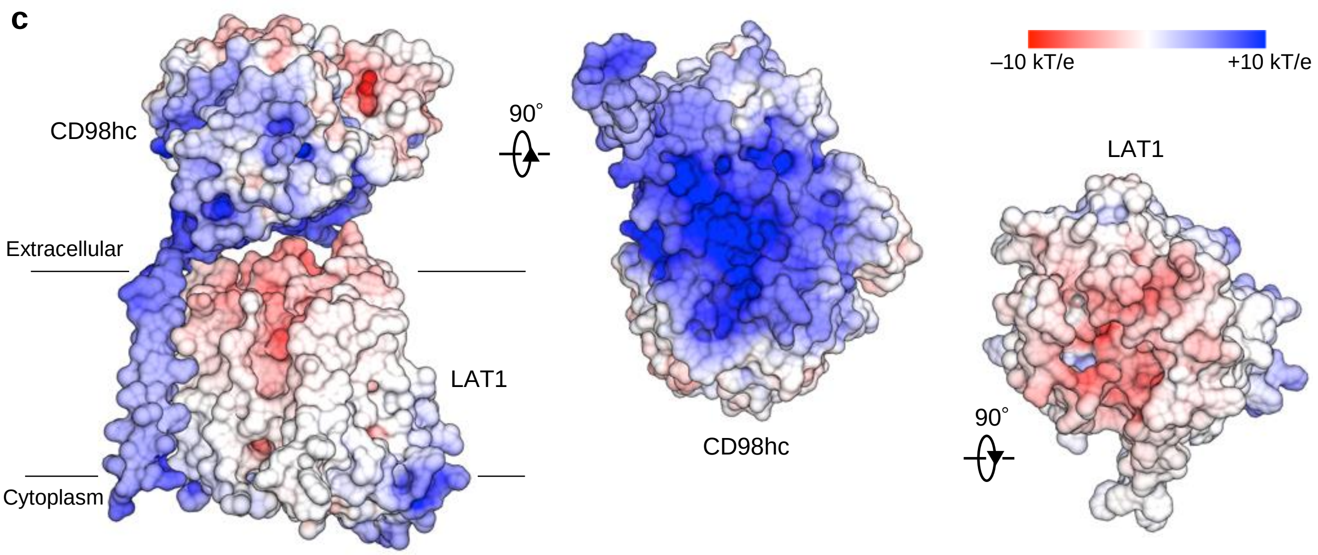

**Figure S5 | Continued**

c) Surface electric potential calculation of LAT1 and CD98hc.

Lee et al. Figure S5 (continued)

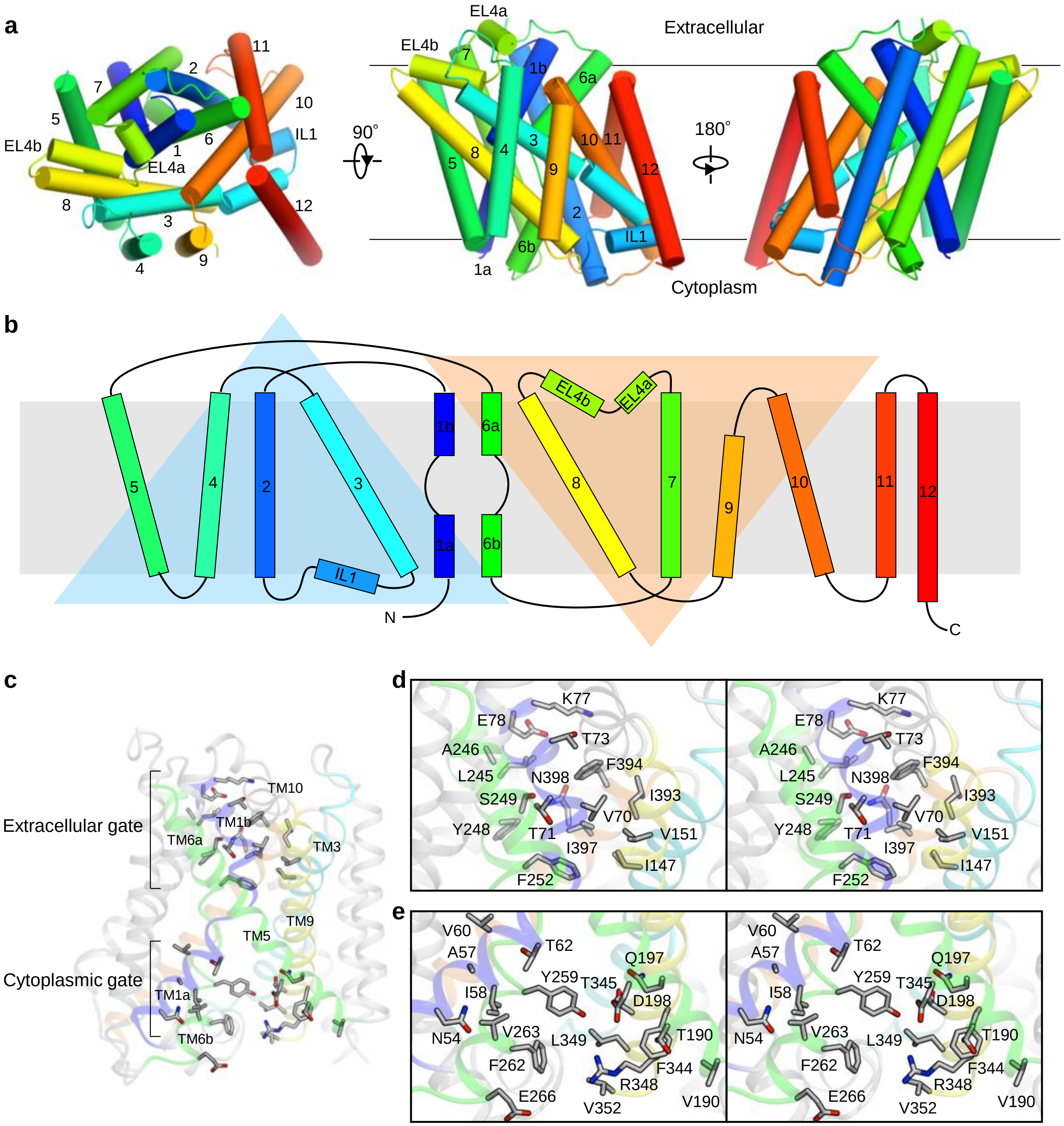

**Figure S6 | Topology of LAT1 and its inward-open state.**

a) Overall structure of LAT1. Helices are colored from blue to red. The structure was visualized from three different viewpoints.
b) Topology diagram of LAT1. The two large triangles indicate the ‘5+5’ inverted repeats.
c) Extracellular and cytoplasmic gates.
d) Stereo view of the extracellular gate.
e) Stereo view of the cytoplasmic gate.

### Lee et al. Figure S6

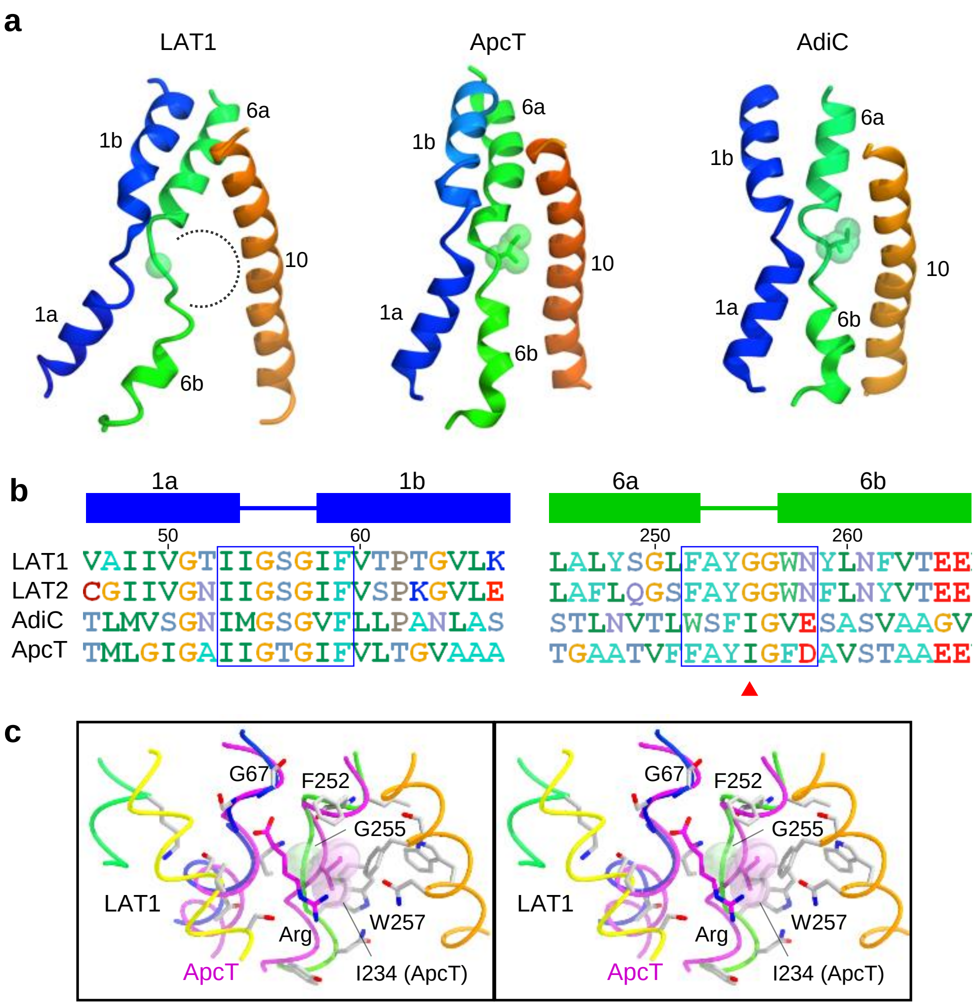

**Figure S7 | Unique configuration of the TM6a-TM6b loop underlies the specificity of LAT1.**

a) Comparison of TM1-TM6-TM10 between LAT1, LAT2, ApcT and AdiC.

b) Sequence comparison of TM1-TM6 in LAT1, ApcT and AdiC. The position of Gly255 is indicated by an red arrow. Blue boxes indicate the loop between the two discontinuous helices.

c) Stereo view of the substrate-binding site of LAT1. The structure of ApcT with bound Arg is superimposed and shown in magenta.

Lee et al. Figure S7

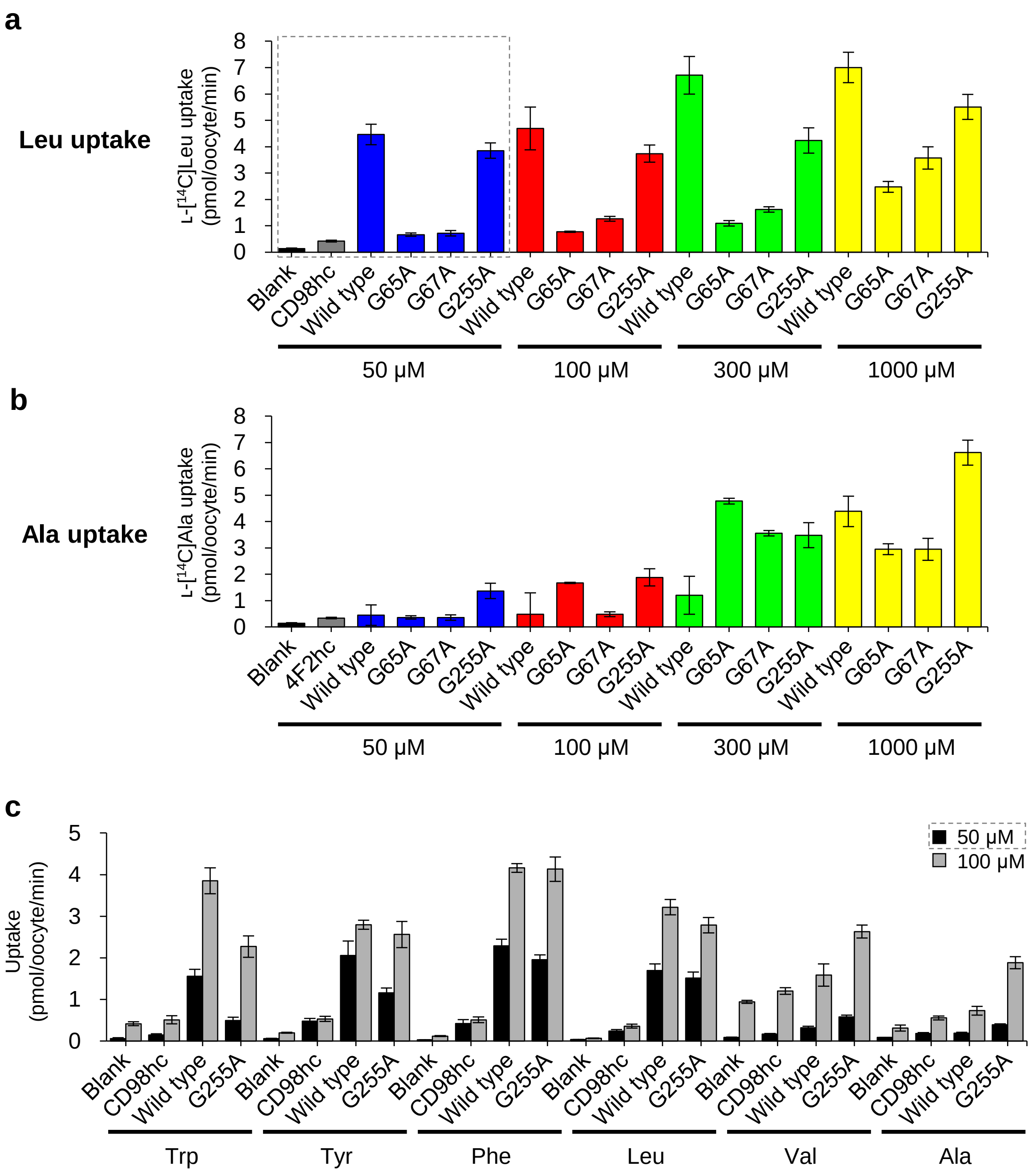

**Figure S8 | Transport assays using *X. laevis* oocytes**

a) Uptake of L-[$^{14}$C]Leu by *X. laevis* oocytes co-expressing CD98hc and LAT1 variants, measured at four different substrate concentrations. The data used for Fig. 4e are marked by gray dotted rectangles.

b) Uptake of L-[$^{14}$C]Ala by *X. laevis* oocytes co-expressing CD98hc and LAT1 variants, measured at four different substrate concentrations.

c) Uptake of various radiolabeled amino acids by *X. laevis* oocytes co-expressing CD98hc and LAT1 (wild type or G255A) measured at two different substrate concentrations. The data from the 50 µM substrate concentrations are used for Fig. 4f.

(Continues on the next page)

Lee et al. Figure S8

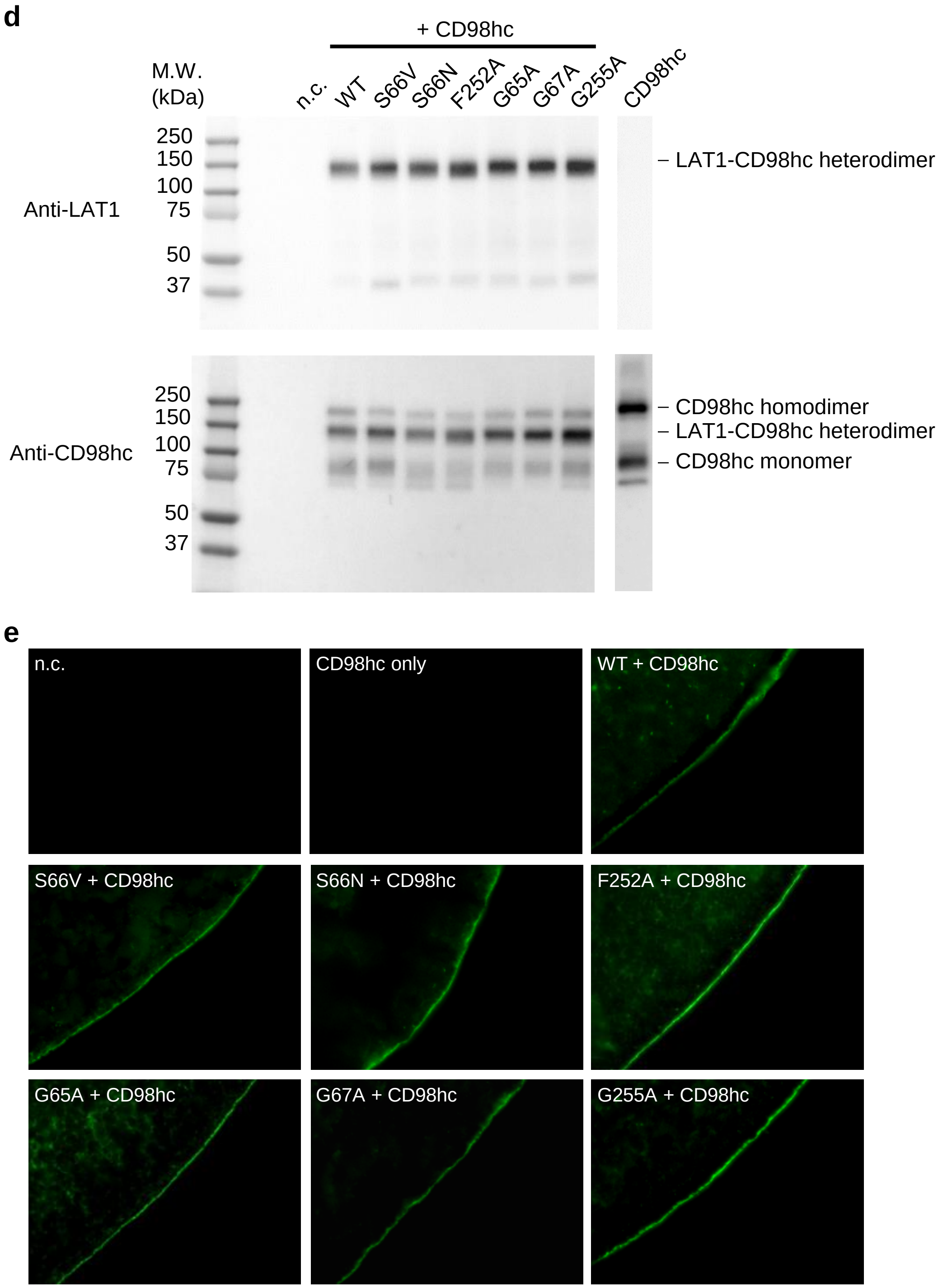

**Figure S8 | continued**

d) Western blotting of CD98hc and LAT1 variants expressed in *X. laevis* oocytes. Isolated oocyte membranes were fractionated by SDS-PAGE in the absence of DTT, and blotted with anti-LAT1 or anti-CD98hc antibodies. The results confirmed the expression and heterodimer formation for all variants.

e) Immunofluorescence of *X. laevis* oocytes, detected by an anti-LAT1 antibody. The results confirmed the plasma membrane localization for all variants.

### Lee et al. Figure S8 (continued)

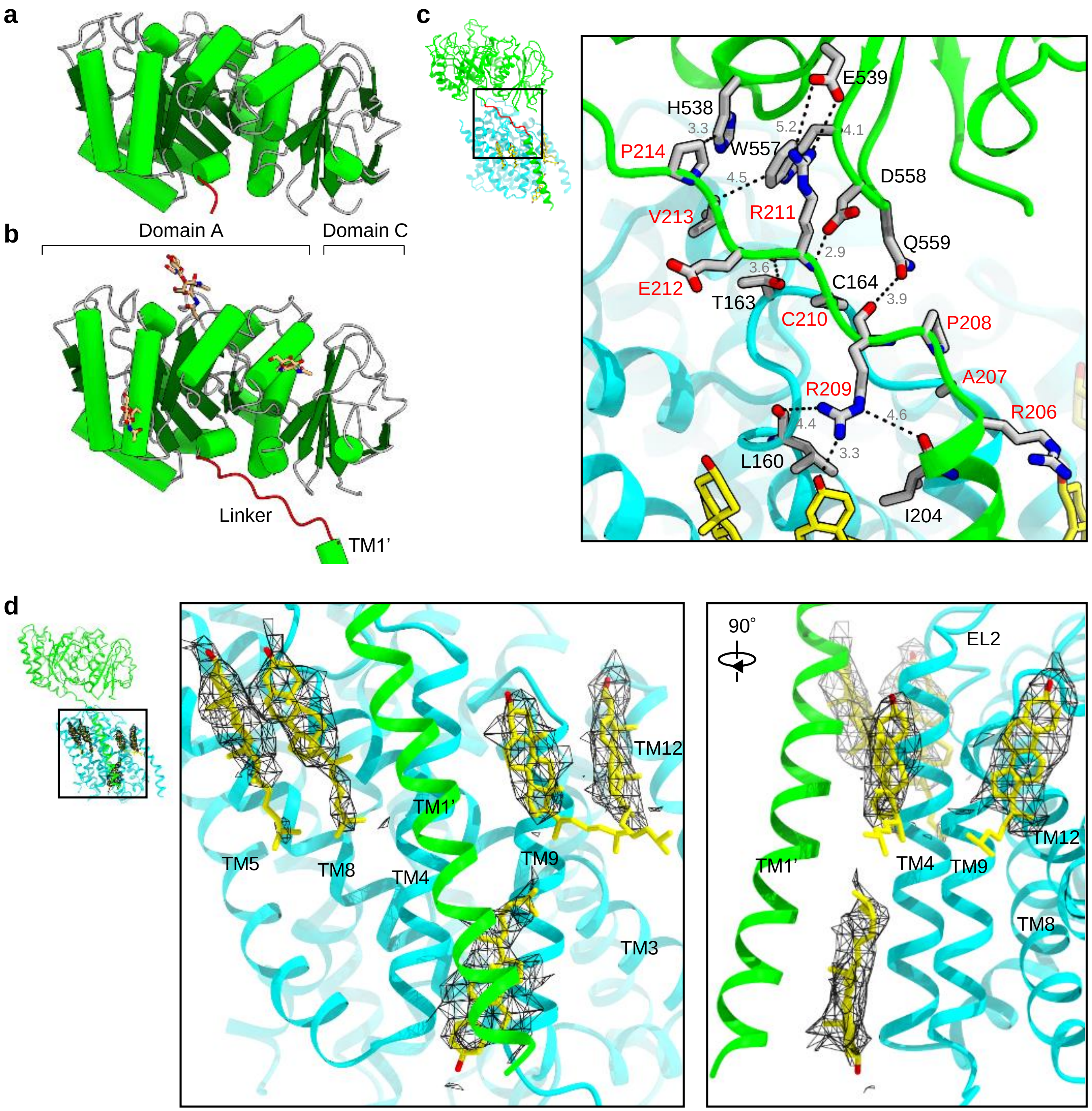

**Figure S9 | Full-length CD98hc and lipid interactions**

a) Crystal structure of CD98hc-ED (PDB: 2DH3).
b) Cryo-EM structure of CD98hc in the LAT1-CD98hc heterodimer.
c) Close-up view of the linker. Atom-atom distances are labeled in gray. Linker residues are labeled in red.
d) Close-up view of the sterol binding sites. Densities are contoured at 11 σ.

Lee et al. Figure S9

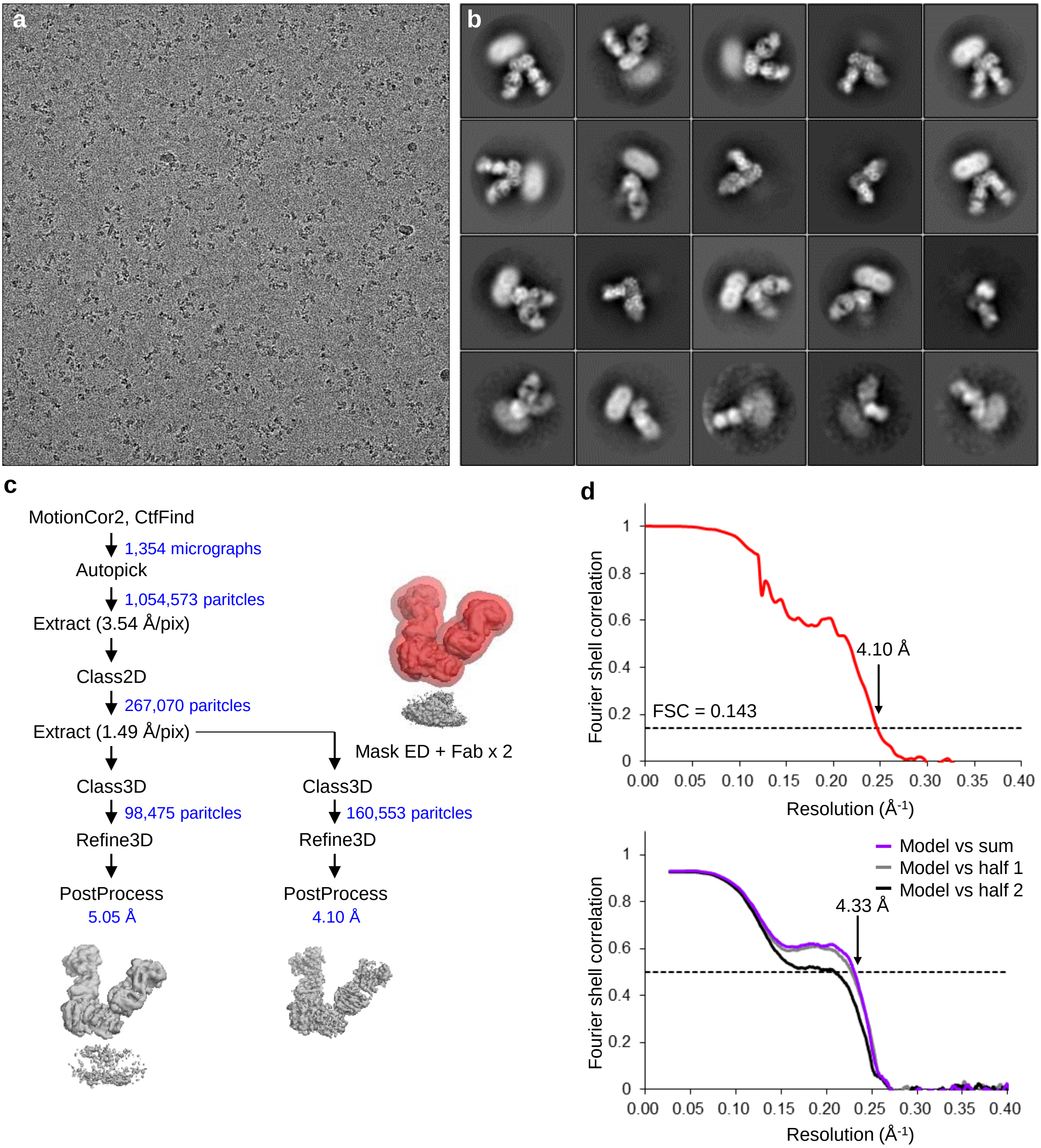

**Figure S10 | High-resolution cryo-EM of LAT1-CD98hc-HBJ127 Fab-MEM-108 Fab**

a) Representative cryo-EM image of LAT1-CD98hc-HBJ127 Fab-MEM-108 Fab, recorded on a 200 kV Tecnai Arctica with a K2 camera.

b) 2D class averages from **a**.

c) Workflow of single-particle image processing.

d) Fourier shell correlation curve of two half-maps.

e) Map-to-model correlation curve.

Lee et al. Figure S10

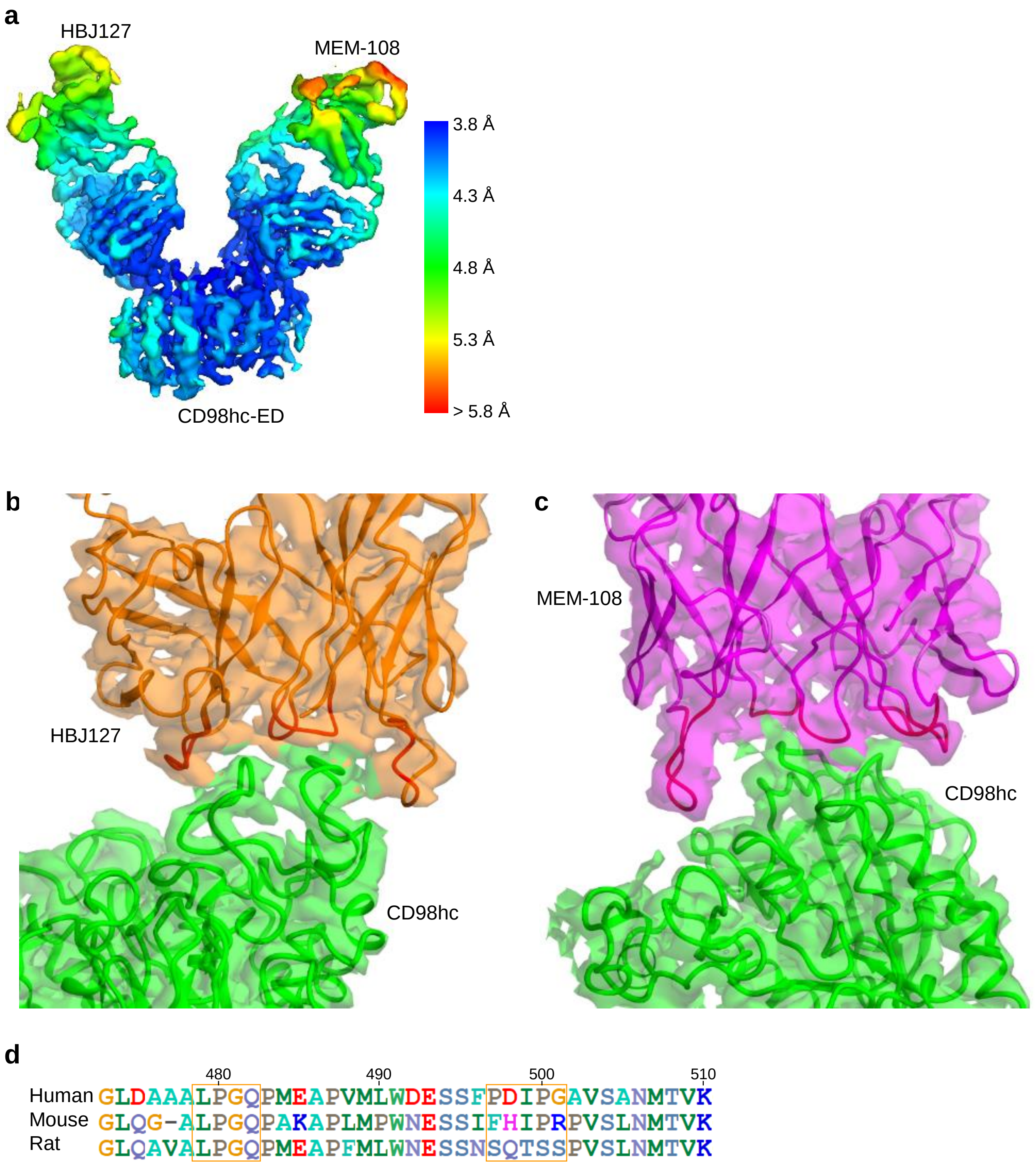

**Figure S11 | Structure of CD98hc-ED-HBJ127 Fab-MEM-108 Fab.**

a) Local resolution analysis.
b) Model of HBJ127 Fab in the density map
c) Model of MEM-108 Fab in the density map.
d) Sequence alignment between human, mouse and rat CD98hc. The HBJ127 epitopes are indicated by orange boxes.

Lee et al. Figure S11

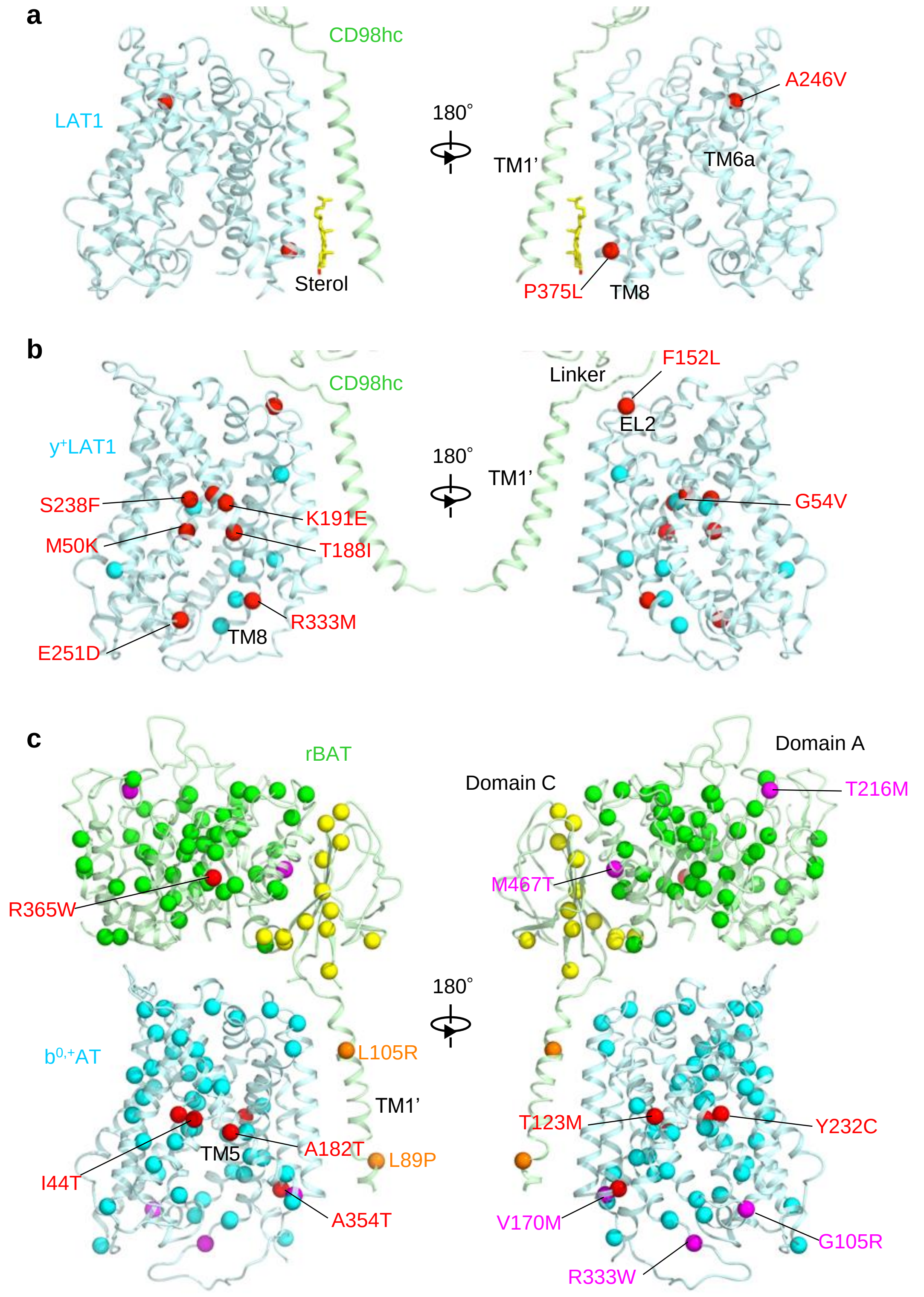

**Figure S12 | Disease-associated mutations of SLC3 and SLC7.**

a) Disease mutations of LAT1. Selected mutations are colored red.

b) Disease mutations of y$^{+}$LAT1.

c) Disease mutations of b$^{0,+}$AT and rBAT. Mutations in TM1’ are colored in orange. Frequent mutations are colored magenta.

Lee et al. Figure S12

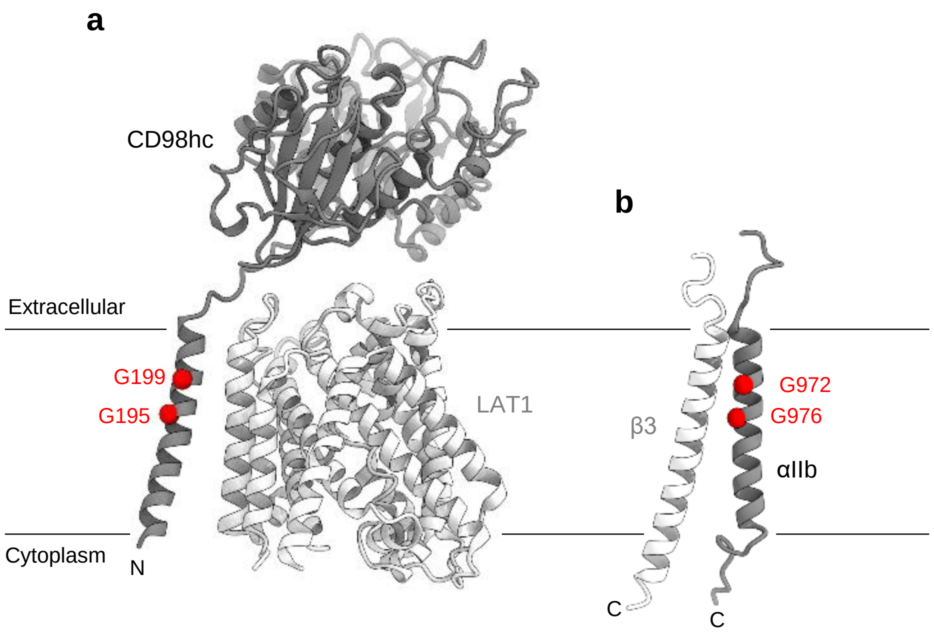

**Figure S13 | GxxxG motif.**

a) GxxxG motif in the LAT1-CD98hc complex. The motif is exposed to the lipid bilayer, and is located roughly at the same depth from the membrane surface as the GxxxG motif in integrin αIIb, which is shown in **b**. It suggests that the GxxxG motif in CD98hc may provide a similar interaction interface for integrin β3.

b) GxxxG motif in the integrin αIIb-β3 complex (PDB ID: 2K9J).

Lee et al. Figure S13

**Table S1 | Cryo-EM data collection, refinement and validation statistics**

| | LAT1-CD98hc-MEM-108 Fab | LAT1-CD98hc-MEM-108 Fab-HBJ127 Fab |
|---|---|---|
| **Data collection and processing** | | |
| Microscope | Titan Krios G3i | Tecnai Arctica |
| Detector | Falcon III | K2 Summit |
| Magnification | 96,000 | 23,500 |
| Voltage (kV) | 300 | 200 |
| Electron exposure ($e^-$/Å$^2$) | 46.0 | 50.2 |
| Defocus range (μm) | –1.5 to –2.5 | –0.5 to –1.5 |
| Pixel size (Å) | 0.861 | 1.490 |
| Symmetry imposed | C1 | C1 |
| Initial particle images (no.) | 635,921 | 1,054,573 |
| Final particle images (no.) | 250,712 | 160,553 |
| Map resolution (Å) | 3.41 | 4.10 |
| FSC threshold | 0.143 | 0.143 |
| Map resolution range | 3.2–5.2 | 3.8–5.8 |
| **Model building and refinement** | | |
| Initial model used (PDB) | 2DH3, 4ZXB | 2DH3, 4ZXB |
| Model Resolution (Å) | 4.02 | 4.33 |
| FSC threshold | 0.5 | 0.5 |
| Map sharpening B factor (Å$^2$) | –30.6 | –145.0 |
| Model composition | | |
| Non-hydrogen atoms | 9,954 | 9,809 |
| Protein residues | 1,282 | 1,287 |
| Ligands[a] | 210 | 70 |
| R.m.s. deviations | | |
| Bond lengths (Å) | 0.005 | 0.010 |
| Bond angles (°) | 0.960 | 1.126 |
| **Validation** | | |
| MolProbity score | 1.46 | 1.89 |
| Clashscore | 2.56 | 6.43 |
| Poor rotamers (%) | 0.29 | 0.84 |
| Ramashandran plot | | |
| Favored (%) | 93.64 | 90.42 |
| Allowed (%) | 6.28 | 0.58 |
| Disallowed (%) | 0.08 | 0.00 |

[a]Ligands include cholesterol and GlcNAc.

Lee et al. Table S1
